# Big Bones Mean Big Muscles: AI Quantifies 71 Individual Muscles Across the Body, Revealing Widespread Links Between Muscle, Bone, and Size

**DOI:** 10.1101/2025.04.01.646599

**Authors:** Lara Riem, Megan Pinette, Olivia DuCharme, Valeria Pabon, Jacob Morris, Ashley Coggins, Liza Harold, Kathryn Eve Costanzo, Matthew Cousins, Raina Hein, Matt Rhodes, Eline Lievens, Xue Feng, Savannah D. Benusa, Tim Breeding, Wim Derave, Silvia S. Blemker

## Abstract

Body sizes and shapes vary widely, even among healthy adults, resulting in diverse muscle sizes, strengths, and performance capacities. This study developed a novel AI-driven algorithm to segment and analyze individual muscles and bones from whole-body MRI. Validated via inter- and intra-observer analyses, the algorithm created 3D segmentations of 71 muscles and 13 bones across the upper limbs, trunk, and lower limbs in 48 healthy adults (24 males, 24 females) aged 18–49 years. Muscle volume, asymmetry, length, and fat fraction were quantified. While asymmetry and fat fraction varied across muscles, they did not differ significantly between sexes. Total muscle volume was the strongest predictor of individual muscle volume, followed by bone volume, which correlated with muscle size at whole-body, regional, and individual levels. Muscle-to-body size relationships (e.g., mass, height, BMI) differed between sexes, while bone-to-body size relationships did not. These findings suggest that skeletal size (“frame size”) is a key determinant of muscularity. This study provides the most comprehensive in vivo dataset of human skeletal muscle to date, offering a multi-factorial explanation for variation in muscularity and benchmarks for applications ranging from athletic performance to clinical assessments in muscle disorders.

**SUMMARY:** Understanding how muscle size varies across the body and what influences these differences is key to advancing both health and performance. This study uses cutting-edge AI to analyze individual muscles across the whole body from MRI scans, creating the most detailed dataset of human muscle and bone relationships to date. The findings reveal how skeletal size (“frame size”) and body dimensions shape muscularity, providing new insights into why people differ in muscle size and strength. These results have broad implications, from optimizing athletic training to diagnosing and treating muscle disorders. By uncovering the intricate connections between muscles, bones, and body size, this research offers a powerful framework for understanding human movement and variability.

## INTRODUCTION

Skeletal muscles are the primary engines powering all human movement. From enabling humans to achieve physical triumphs such as breaking Olympic weight-lifting or sprinting records to providing the fine motor control required for piano playing or performing surgical procedures, sufficiently developed muscles are essential for much of human behavior. This vast range of functions is achieved through variation in size, shape, and architecture of skeletal muscles across muscles in the human body. Humans display a remarkable range in body size, with most adults falling between 5 and 7 feet in height and between 100 and 300 pounds in weight. Just as the shapes and sizes vary across different muscles, the shapes and sizes of each muscle vary significantly across people. What determines individual muscularity, and why are some people naturally more muscular than others? The factors influencing muscle size variation—referred to as ’scaling relationships’—across different body shapes and sizes are not fully understood.

Previous studies exploring scaling relationships have examined muscles and bones across species of varying body sizes, often using body mass as a predictor of muscle size.^1,2^ In humans, studies have examined muscle shape and sizes at the individual muscle level,^3^ muscle group level,^4^ and at the whole upper or lower extremity level.^5,6^ These studies demonstrated how muscle size varies across individuals with different sexes, heights, masses, BMIs, performance ability, and neuromuscular and musculoskeletal conditions.^7^ The wide variation in muscle size across populations demonstrates high adaptability of muscle to power movement and the susceptibility of muscle to disuse and disease across a range of individuals. This variability in muscle sizes directly influences functional capabilities, from strength^8^ to motor control.^9^ Across general healthy population, muscles generally scale together within specific regions. For example, shoulder, elbow, and wrist muscle volumes maintained percent relative to total upper extremity muscle volume across individuals^6^. A similar finding has been seen in muscle groups in the lower extremity.^5^ Furthermore, scaling relationships between muscle volumes and the product of height and mass illustrate the novel concept of muscle scaling within the human population. Most of these relationships are considered functionally intuitive given that these muscles have anatomical proximity and are functionally synergistic. When muscles are not anatomically close or part of the same functional groups, do they still scale with one another? Can muscle size variations across the entire body be explained by differences in height and mass, as observed in lower limb studies?^5^ How do these variations differ between males and females?

Characterizing the principles that govern muscle variation in healthy populations has wide-reaching potential. Understanding how individual muscle sizes scale can help establish expected values for specific muscles in a given person. Assessing deviations from these expected muscle sizes can provide insights into musculoskeletal health and performance across various patient and athlete populations. Data on muscle scaling can lead to more representative musculoskeletal models, with applications ranging from robotics to surgical planning. Finally, characterizing muscle diversity among individuals will enhance our understanding of healthy body composition across various body sizes and shapes.

Given the complexity and variability in muscle anatomy, accurate and efficient segmentation is crucial for understanding these scaling relationships. Segmentation of muscles from magnetic resonance imaging (MRI) is the most advanced and precise method for quantifying muscle volumes *in vivo*; however, muscle segmentation is a notoriously challenging and time-intensive task due to the complex anatomy of many muscles, the subtle boundaries between muscles, and variable anatomy of muscles across different individuals. Manual segmentation, which has historically been the standard, is not only labor-intensive but also prone to variability depending on the operator’s experience and the quality of imaging data. To address these challenges, we have pioneered an AI-driven convolutional neural network (CNN) approach to muscle segmentation, beginning with the healthy lower limb muscles,^10^ muscles in patients with neuromuscular disease,^11^ and rotator cuff muscles.^12^ Our method leverages deep learning techniques to automate the segmentation process, significantly reducing the time required and improving the consistency of results. While other studies have also confirmed the utility of CNNs for muscle segmentation, no one to date has developed an automatic segmentation approach for the whole body.

The goals of this work are to: (i) develop a CNN algorithm to provide an AI-powered full body muscle segmentation method; (ii) use the method to quantify the volumes, lengths, and fat infiltration of 71 unique muscles and 13 unique bones across the upper extremity, trunk, and lower extremity in a group of healthy individuals ranging in age from 18 to 49; and (iii) examine muscle scaling relationships across individuals of varying heights and weights, with attention to characterizing sex differences. These measurements will provide a detailed characterization of muscle architecture, essential for understanding how muscle size and shape vary across different body sizes and compositions. This work will not only advance our understanding of human musculoskeletal design but also enhance the accuracy of musculoskeletal modeling, with broad implications for a wide range of health applications.

## RESULTS

### Volume fraction, length, asymmetry, and fat fraction across muscles

Generally, the distribution of muscles **(Fig. 2)** across the body is preserved across the population studied: standard deviations of muscle volume percentage varied from 2% to less than 0.1%. The largest muscles, as measured by percent of total volume, were found in the lower limb (gluteus maximus – average of 7.8% in males and 8.7% in females); while the smallest muscles were found in the upper limb (primarily wrist muscles, average of 0.07% in both males and female). The highest levels of asymmetry (**Fig. 3A**) were observed in the upper extremity muscles (wrist flexor and extensor muscles). When expressed as a percentage of the associated bone length, muscle lengths (**Table 2**) were highest in the upper limb (trapezius and multifidus) and lowest in the lower limb. The highest average fat infiltration levels (**Fig. 3B**) were also seen in upper limb muscles (scapular and trunk muscles). Female and male muscles were most different in muscle volume percentage (higher lower extremity muscle volume percentages in females; higher upper limb muscle volume percentages in males) and fat (female muscles often had slightly higher fat levels than male muscles). There were no significant sex differences in muscle length to bone length scaling relationships (**Supplemental Materials**). All descriptive results, main effects, interaction effects, and post-hoc analysis results from the asymmetry, fat infiltration, normalized volume, volume fraction, and difference between actual and predicted volume two-way mixed repeated ANOVA statistical analyses are reported in **Supplemental Material Tables S3-S8.**

**Figure 1:**
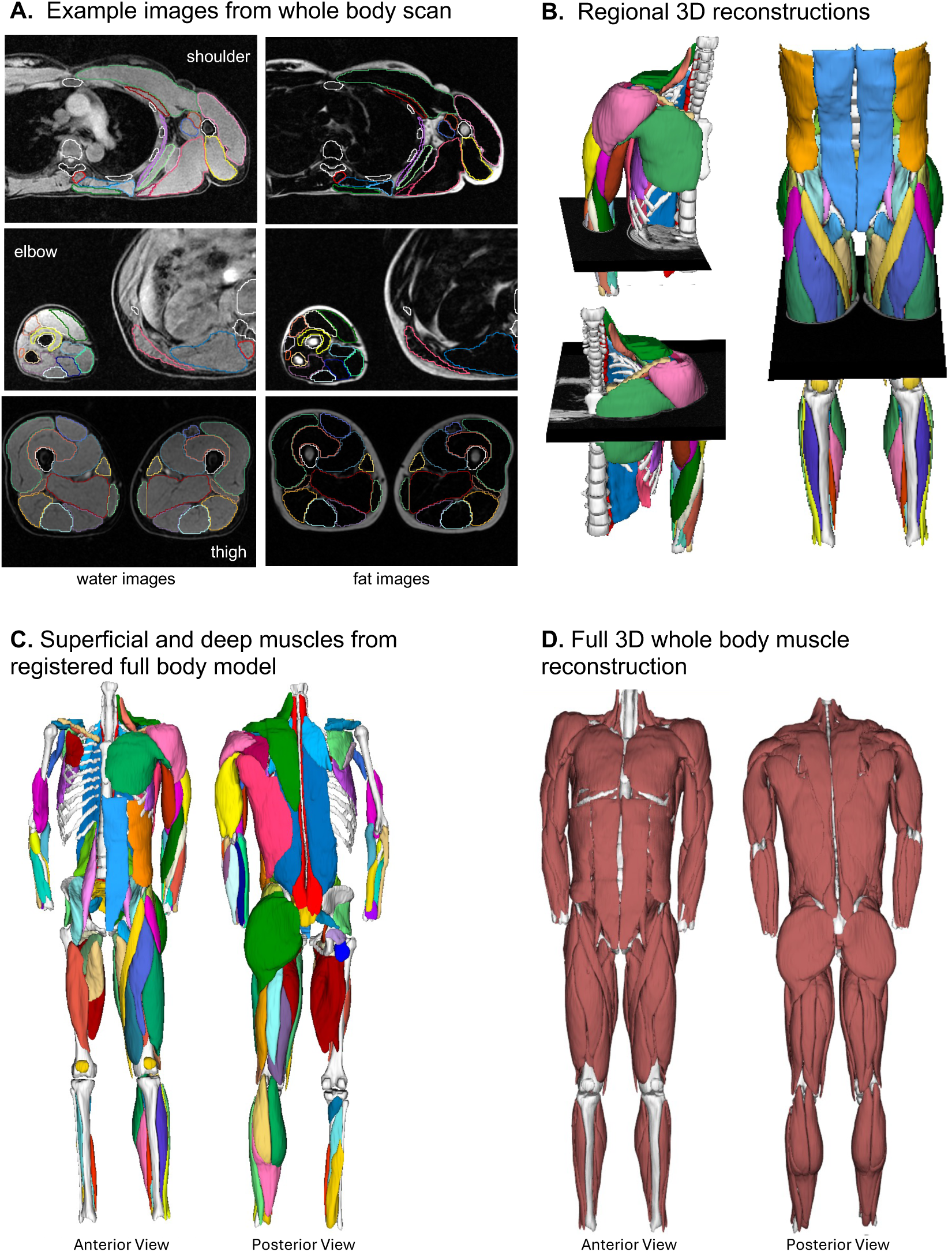
Whole-body imaging and muscle segmentation. (A) Sample axial slices of fat and water images from the shoulder, elbow, and thigh regions. (B) Regional 3D reconstructions for each segmented body region (UBL, UBR, LE_EXT), shown with corresponding axial slices. (C) Full-body reconstruction displaying superficial muscles on the left anatomical side and deep muscles on the right side. (D) Registered full-body reconstruction in the native anatomical view.

**Figure 2.**
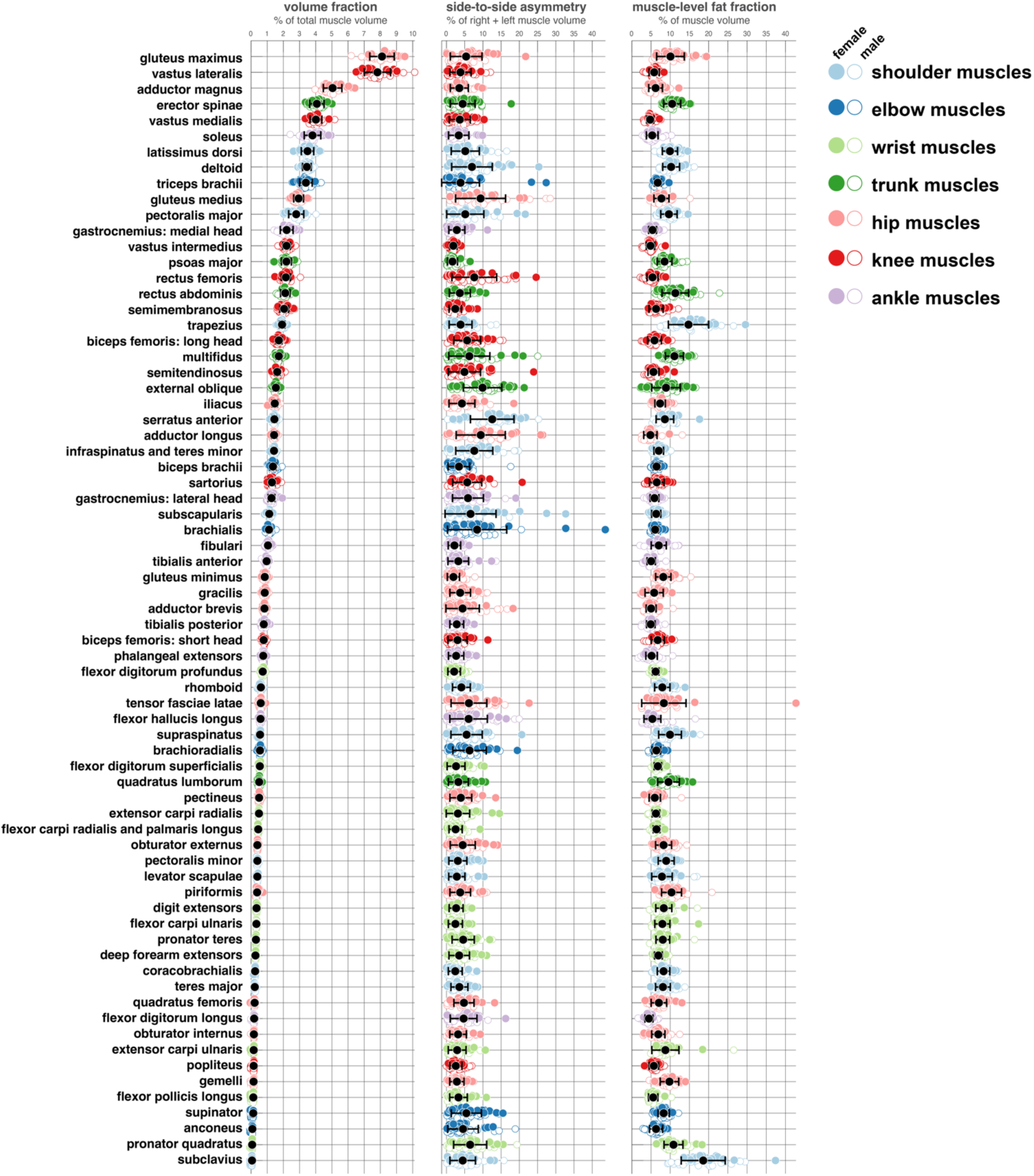
Comparison of volume fraction, right-left asymmetry, and fat fraction across 71 muscles in the human body. Muscles are ordered by average volume fraction, from largest (top) to smallest (bottom). High and low levels of asymmetry and fat fraction are present across the body, with higher values generally observed in upper-body muscles (shoulder, elbow, wrist, trunk) compared to lower-body muscles (hip, knee, ankle). Analysis of differences in these measures are provided in Supplemental Data.

**Figure 3.**
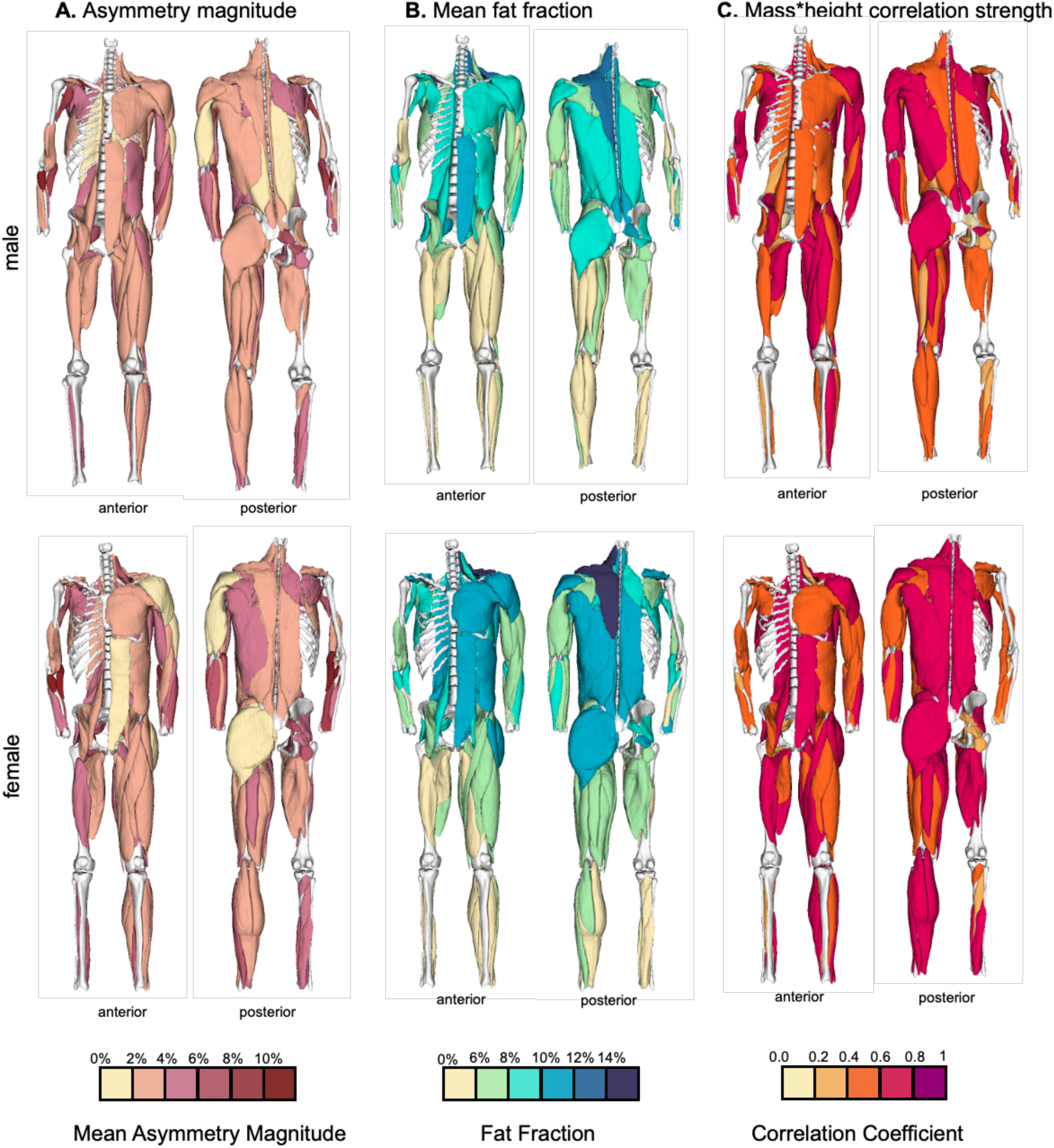
Average asymmetry magnitude, fat fraction percentage, and correlation coefficients for male and female cohorts mapped onto sample muscle models. Top Left: Male asymmetry magnitude values shown in superficial (left) and deep (right) views. Bottom Left: Female asymmetry magnitude values shown in superficial (right) and deep (left) views. Top Middle: Male fat fraction values shown in superficial (left) and deep (right) views. Bottom Middle: Female fat fraction values shown in superficial (right) and deep (left) views. Top Right: Male correlation coefficients for the height*mass method shown in superficial (right) and deep (left) views. Bottom Right: Female correlation coefficients for the height*mass method shown in superficial (right) and deep (left) views.

**Table 2.**
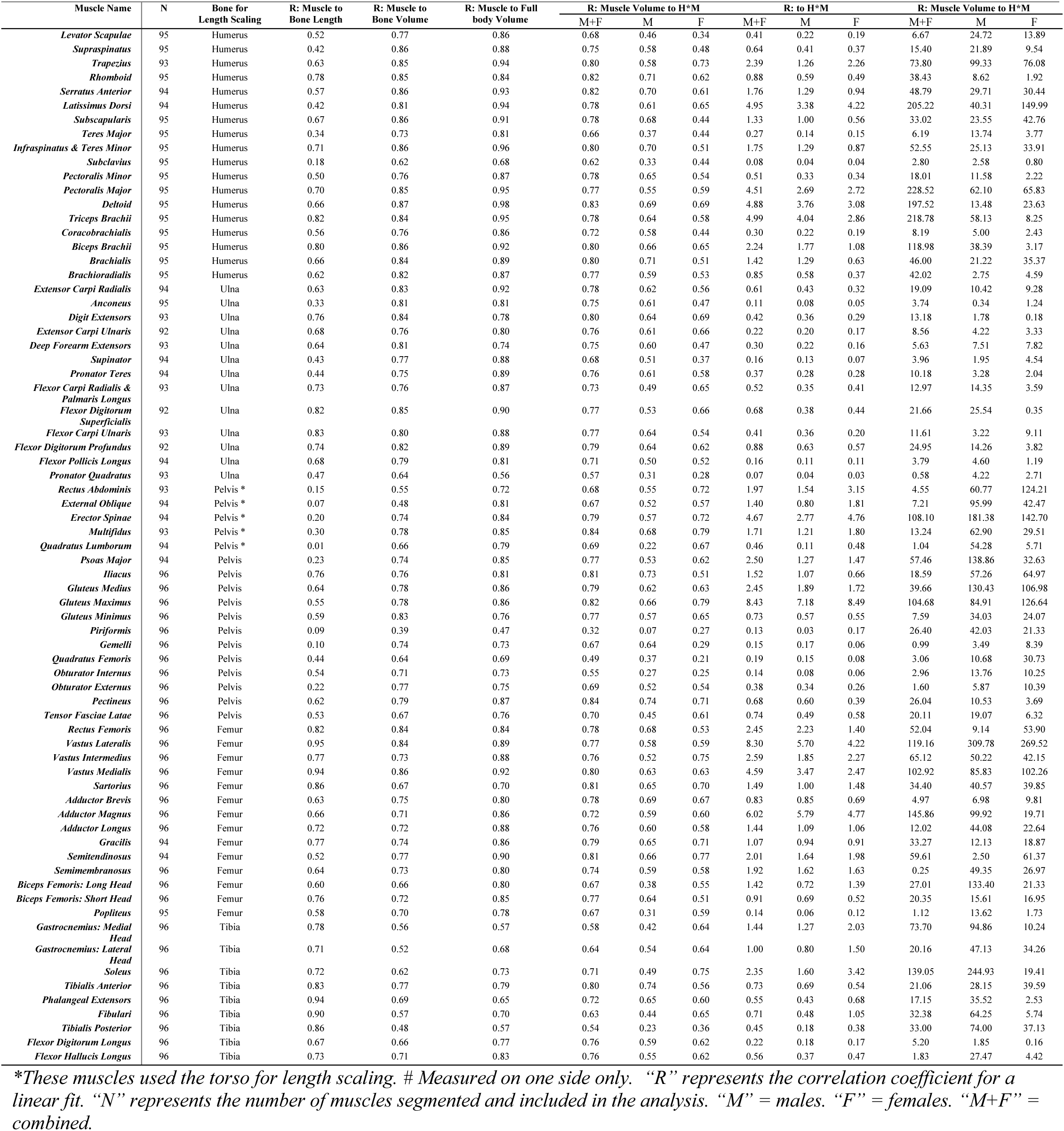
Scaling relationships between bone length, volume, body size, muscle length, and muscle volume for each individual muscle.

### Individual muscle volumes are best predicted by total muscle volume and associated bone volume

Across upper and lower limb muscles, individual muscle volumes **(Fig. 4**, **Table 2)** demonstrated the highest correlation coefficients with total muscle volume, followed by associated bone volume, mass*height, mass, height, and then BMI, with a few exceptions. First, a selection of very small muscles had similarly low correlations across all measures (subclavius, pronator quadratus, piriformis, tibialis posterior). Second, several wrist extensor muscles scaled most strongly with associated bone volume, while several ankle dorsiflexor muscles and the sartorius scaled most strongly with body dimensions (mass*height or mass). Of the measures that do not require quantification of other muscles or bones, product of height and mass scaled most strongly with individual muscle volumes (**Fig. 4**, **Fig. 3C**). The muscles crossing the shoulder, elbow, and knee joint had the highest correlation coefficients across scaling parameters; by contrast, the muscles crossing the ankle joint had lowest coefficients. While correlation coefficients differed across parameters, there was no statistically significant main effect of muscle or method, but there were significant interaction effects (**Supplemental Material Table S2**).

**Figure 4.**
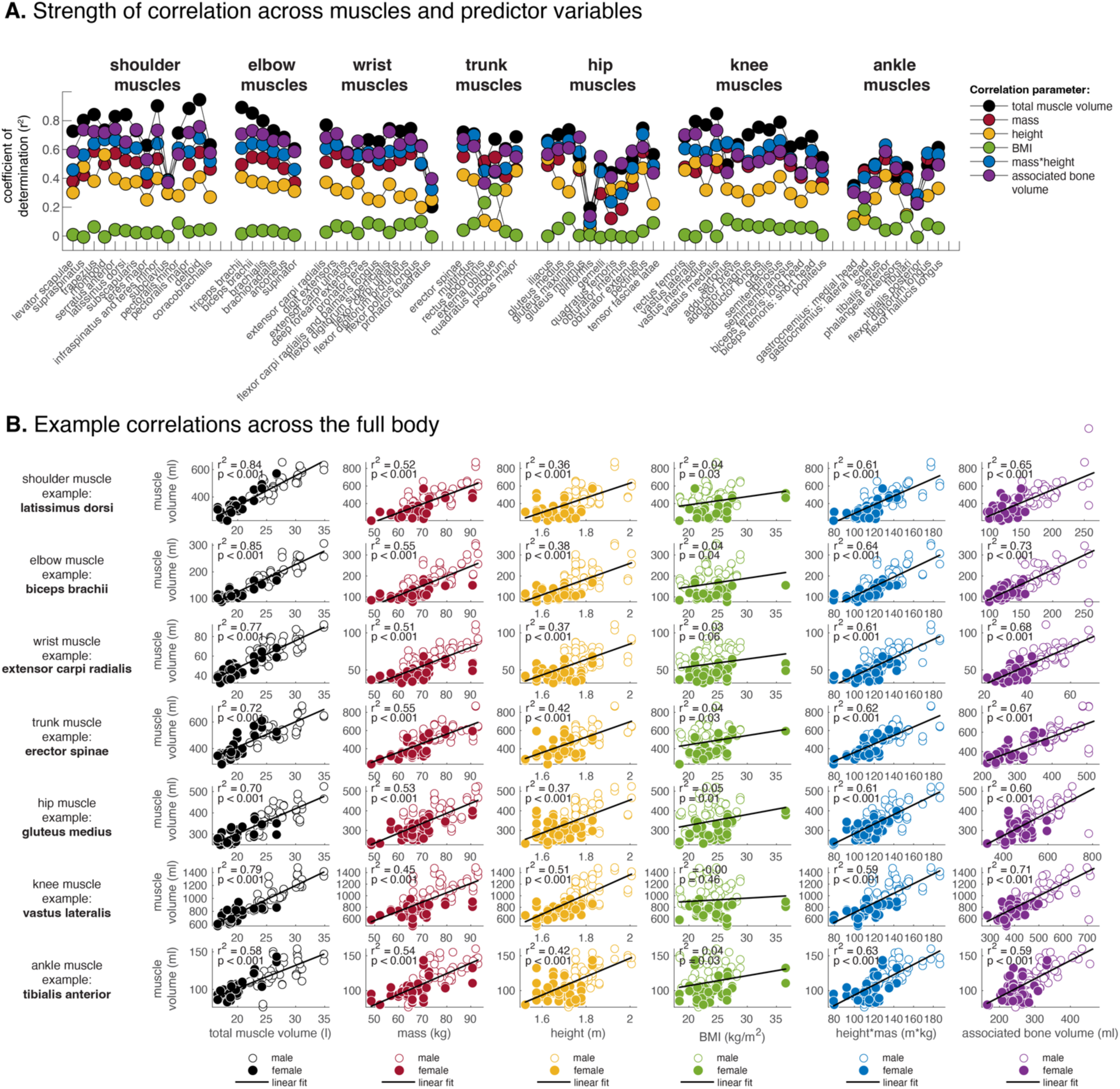
Correlation of individual muscle volumes with body parameters, associated bone size, and total muscle volume across 71 measured muscles. Total muscle volume was the strongest predictor, followed by associated bone volume, height*mass, body mass, height, and BMI

### Muscle distribution patterns are unique and different between female and male subjects

Across subjects, all but three female subjects and one male subject clustered (**Fig. 5**) with other subjects of the same sex. Trunk muscles, calf muscles, deep hip muscles (piriformis and gluteus minimus) and one hamstring muscle (semimembranosus) had a pattern of higher volume fraction amongst the cluster on the left of the matrix that contained mainly female subjects. By contrast, shoulder, elbow, wrist, and hip muscles had a pattern of higher volume fraction in the cluster on the right of the matrix that contained male subjects. Several other muscles clustered together but did not have a monotonic pattern of difference between male and female subjects.

**Figure 5.**
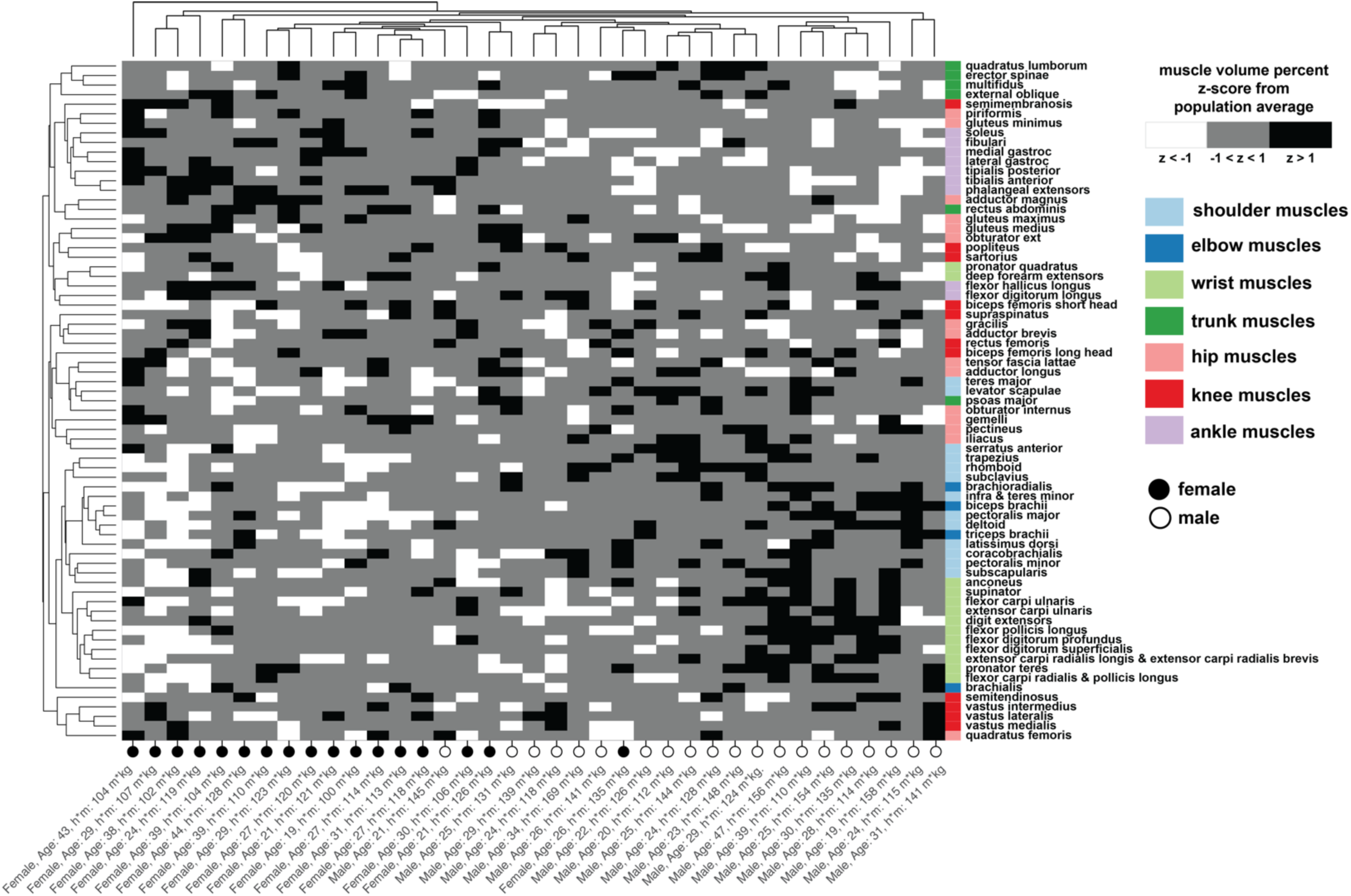
Hierarchical cluster analysis of muscle volume fractions, clustering individuals (columns) and muscles (rows) by similar patterns in muscle volume fraction distribution.

### Upper vs. Lower Body Scaling with Bone Volume

Summed upper body muscle volumes (**Fig. 6A**) correlated strongly with summed lower body muscles; similarly, summed upper body bone volumes correlated strongly with summed lower body muscle volumes. However, the ratio of upper limb to lower limb muscle volume (**Fig. 6B**) was significantly greater in male subjects as compared to female subjects; the ratio of upper body bone volume to lower body bone volume was also significantly greater in male subjects as compared to female subjects. The ratios of upper to lower muscle volumes correlated strongly with the ratio of upper to lower bone volumes. The ratios of muscle volume to bone volume (**Fig. 6C**) were not significantly different between males and females in either the upper or lower limbs. These ratios were, however, strongly correlated with each other.

**Figure 6.**
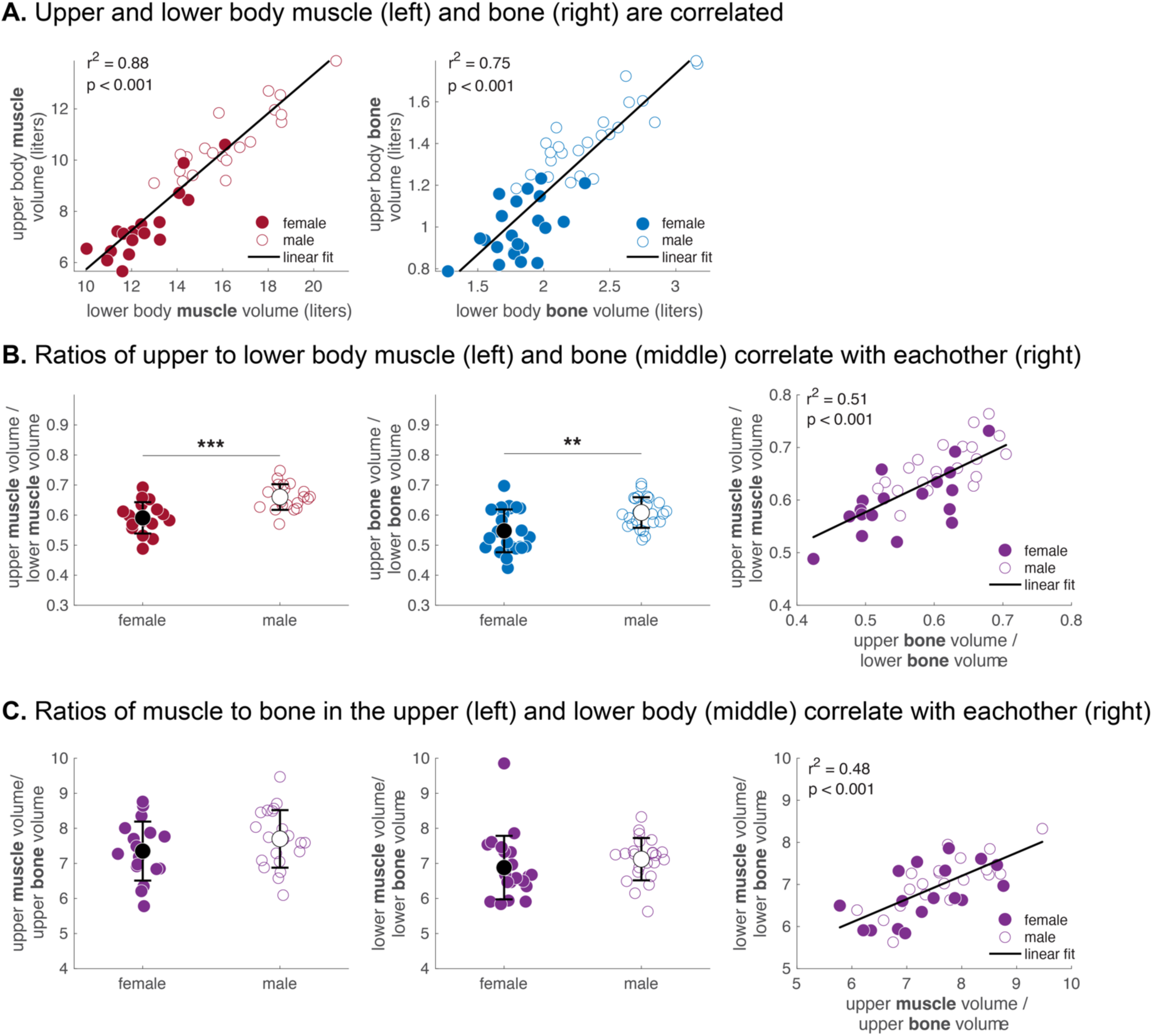
Examination of upper body vs. lower body muscles and bone volumes. (A) Left: Upper body muscle volume vs. lower body muscle volumes. Right: Upper body bone volume vs. lower body bone volumes. (B) Left: Ratios of upper- to lower-body muscle volumes, showing significantly smaller values in females than males. Right: Ratios of upper-to-lower body bone volume, strongly correlating with upper-to-lower body muscle volume ratio. (C) Ratios of muscle volume to bone volume in upper body (left) and lower body (middle), showing no significant differences between sexes. Right: Correlation between upper and lower body muscle-to-bone volume ratios in males and females.

### Total muscle and bone volume scaling

Total muscle volume (**Fig. 7A**) scaled best with total bone volume (r^2^ = 0.76), followed by mass*height (r^2^ = 0.64), mass (r^2^ = 0.52), height (r^2^ = 0.46), and BMI (r^2^ = 0.02). The errors (**Fig. 7B**) between the measured total muscle volume and total muscle volume predicted from each of the five scaling methods were statistically different between female and male subjects for mass (p<0.001), height (p<0.001), BMI (p<0.001), and mass*height (p=0.0012). In all cases, the errors were more negative in female subjects (i.e., scaling prediction overestimated muscle volume) and more positive in male subjects (i.e., scaling prediction underestimated muscle volume), though mass*height predicted volume error had the smallest difference between female and male subjects (-1.7 liters in females vs. +1.4 liters in males). The error between measured and predicted from the bone volume scaling method was not significantly different between female and male subjects. Total bone volume (**Fig. 7C**) scaled best with the product of height*mass (r^2^ = 0.74), followed by height (r^2^ = 0.69), mass (r^2^ = 0.58), and BMI (r^2^ = 0.02). The errors (**Fig. 7D**) between the measured total bone volume and total bone volume predicted from each of the four scaling methods were statistically different between female and male subjects for mass (p<0.001), height (p<0.01), BMI (p<0.001), and mass*height (p<0.01). In all cases, the errors were more negative in female subjects (i.e., scaling prediction overestimated bone volume) and more positive in male subjects (i.e., scaling prediction underestimated bone volume).

**Figure 7.**
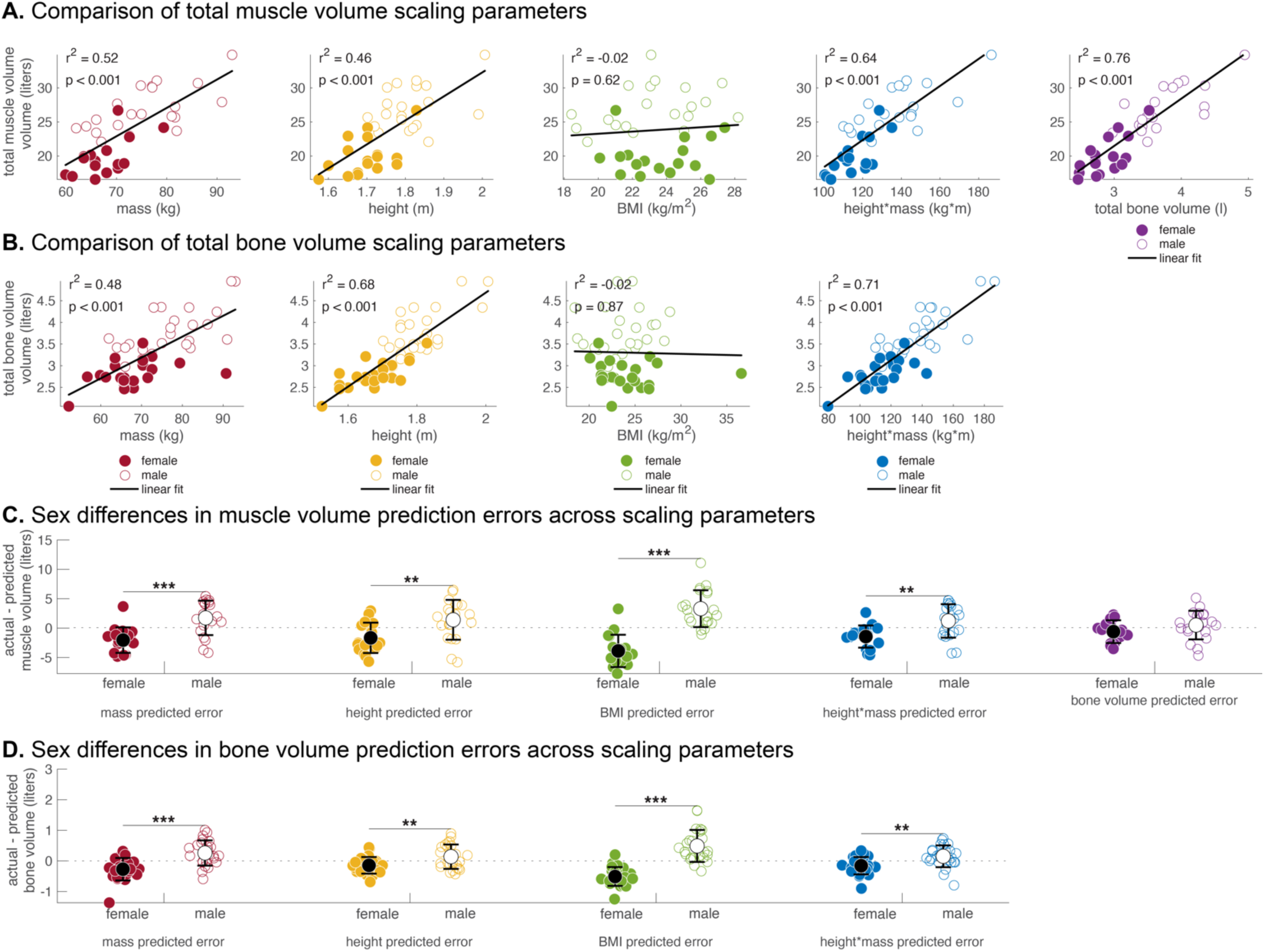
Examination of total muscle volume and total bone volume scaling relationships. Total muscle volume (A) scales most strongly with total bone volume (A-right), followed by height*mass, mass, height, and then BMI. Total bone volume (B) scales most strongly with height*mass (B-right). The error between measured muscle volume and muscle volumed (C) predicted from scaling relationships were significantly less (more negative) in female subjects as compared to male subjects for mass, height, BMI, and height*mass scaling parameters. However, errors between predicted and measured muscle volume did not differ when bone volume was used as the scaling parameter. The error between predicted and measured bone volume (D) differed significantly between female and subjects for all body size parameters (mass, height, BMI, and height*mass).

### Height-bone-muscle combined scaling

Total bone volume per height z-scores accounted for 55% of the variability in total muscle volume per height z-scores (**Fig. 8A**). Similarly, associated bone volume per height z-scores explained 41% of the variability in individual muscle volume per height z-scores (**Fig. 8B**). Overlaid colormaps show that deviations from the predicted muscle and bone z-score relationship (above or below the linear fit line) align with increases or decreases in total muscle volume per total bone volume z-scores (**Fig. 8A**) and in individual muscle volume per associated bone volume (Fig. 8B).

**Figure 8.**
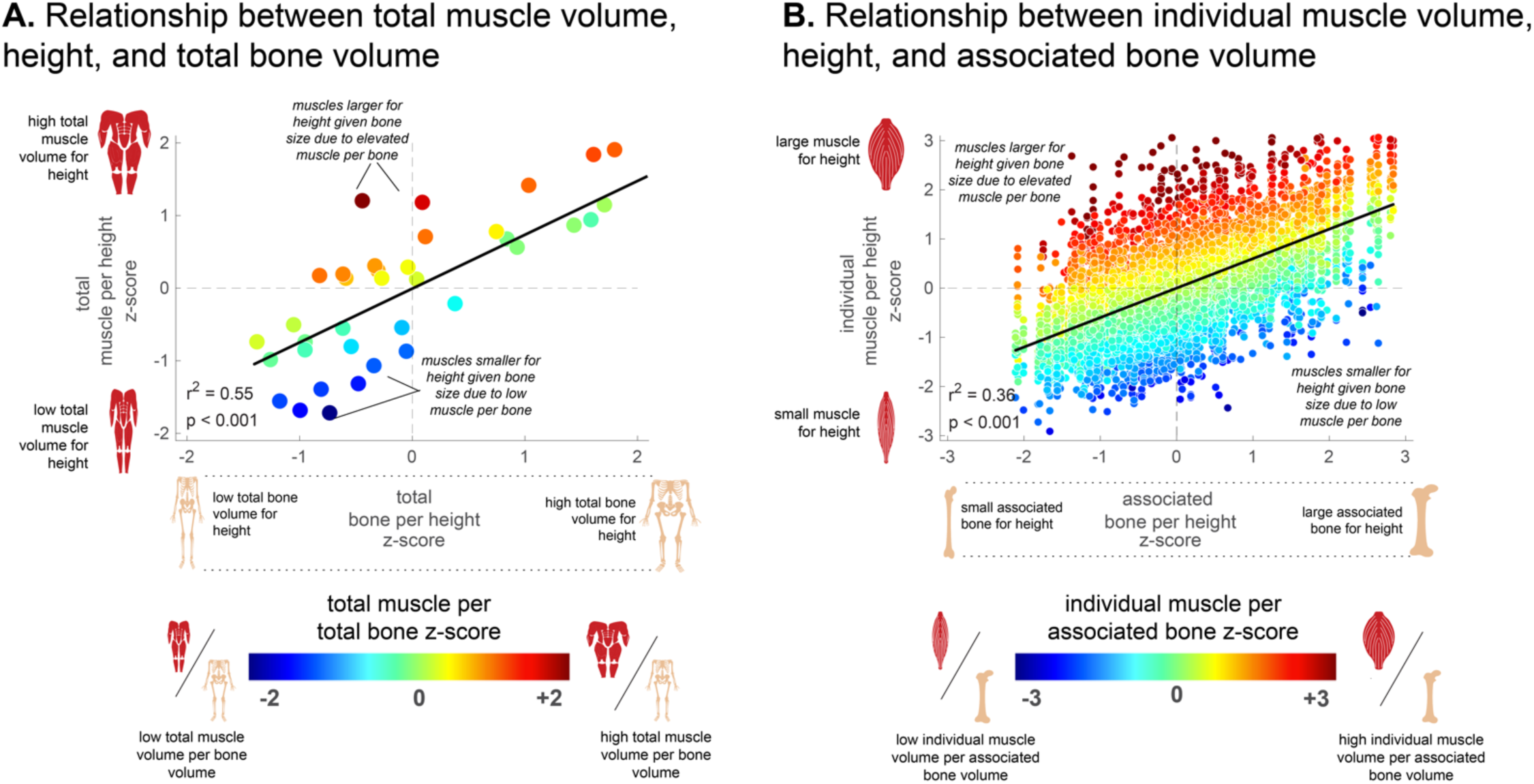
Both total muscle volume scaling (A) and individual muscle scaling (B) can be explained by a combination of height, bone size relative to height and muscle size relative to bone.

## DISCUSSION

The overall goal of this study was to determine which parameters dictate variation in muscle sizes across a cohort of healthy subjects, including muscles of the lower extremity, upper extremity, and trunk bilaterally (71 unique muscles). The study was made possible by the development of a convolutional neural network approach to automatically segment muscles and bones from MR images, which demonstrated high accuracy as compared to intra- and inter-observer variability metrics. This wealth of muscle and bone volume data provided multiple novel insights into muscle variation across individuals, as described in the following paragraphs.

Muscle volume scaled strongly with total bone volume at the individual muscle level (**Fig. 8B**), the body region level (**Fig. 6**), and the total body level (**Fig. 8A**). By expressing muscle and bone volumes as z-scores relative to predicted values based on height, we observed that bone size relative to height (i.e., ‘big bones’ vs. ‘small bones’) strongly correlates with muscle size relative to height (i.e., ‘big muscles’ vs. ‘small muscles’). Prior studies, although not as detailed or comprehensive as this analysis, have similarly found correlations between bone and muscle size.^13,14^ Given that bone size (often referred to as ‘frame size’ or ‘skeletal robustness’) is thought to be largely genetically determined—with some exceptions, such as elite youth athletes or cases of childhood neuromuscular disease—our findings suggest that similar genetic factors may also influence muscle size variation. A genetic basis for co-variation in muscle and bone size has been suggested previously.^14^ Other possible mechanisms of muscle-bone crosstalk, including biochemical, biomechanical, and metabolic factors, have also been demonstrated.^15,16^ These mechanisms, however, primarily affect bone composition parameters, such as cortical thickness, trabecular density, and overall bone mineral density—measures not included in this study. Future work incorporating bone composition analysis would deepen our understanding of the interactions between bone and muscle architectural parameters.

Deviations in the bone z-score to muscle z-score relationship can be explained by the z-score from the muscle volume vs. bone volume relationship (colormap in **Figs. 8A & B**). Specifically, muscle size relative to bone size varied across the population. This variation in muscle size relative to bone size is likely influenced by several factors, such as age, injury, exercise, strength training, nutrition, and overall health status. Together, these observed relationships between muscle, bone, and body size suggest a conceptual framework in which both ‘nature’ and ‘nurture’ factors contribute to an individual’s muscularity and resulting strength.

Of the body size measures explored, individual muscle volumes and total muscle volume scale most strongly with the product of mass and height (mass*height), followed by mass, then height, then BMI. This finding is consistent with our previous work that height*mass was the best predictor of lower limb muscle volumes in twenty-four healthy young adults.^7^ The postulated rationales behind this strong correlation were: (1) there was already a correlation between muscle volume and mass; (2) the addition of height to muscle volume scaling was important for bipeds, suggested by the increased energetic cost and longer muscles of bipeds compared to equal mass quadrupeds;^17^ and (3) the function of muscle is to generate joint torque, which correlates with muscle volume^18^; both mass and height influence the torque required by muscles to provide support and propel the body. It is somewhat surprising that a similar relationship was observed in the upper extremity muscles, even though upper extremity muscles do not support the load of body weight. A fourth explanation could be the observation that bone volume also correlates strongly with height*mass (**Fig. 7C**), which we have already established as an even stronger predictor of muscle volume.

We identified several important differences between male and female subjects, which aligns with prior studies showing significant differences in muscle mass between sexes through measurements at the whole-body level^19^, within the lower limbs^20^, and at the individual muscle level.^21–23^ Skeletal and joint anatomy also varies significantly between males and females.^20,24–27^ Consistent these findings, we observed significant sex differences in scaling relationships between muscle volume and body dimensions: errors between measured and predicted muscle volumes based on height*mass, height, mass, and BMI were significantly more negative in female subjects (and more positive in male subjects). However, a novel aspect of our study is the finding that errors between actual and predicted muscle volume did not differ between males and females when associated bone volume was used as a predictor. This result indicates that after accounting for bone size relative to height, male and female muscle sizes are comparable. This finding suggests that factors regulating bone size relative to height (which tends to be larger in males) may also drive differences in muscle size relative to height between males and females.

The patterns of muscle distribution differed substantially between males and females, as illustrated by muscle the clustering analysis at the muscle-by-muscle level (**Fig. 4**) and the comparison of upper vs. lower body muscle volumes (**Fig. 5**). Generally, female subjects had relatively more muscle in the lower limbs, particularly in the ankle. By contrast, male subjects had more muscle in the upper limbs, particularly in the wrist and shoulder. The investigation of muscle and biological sex-based differences in asymmetry magnitude and fat fraction provided insight about expected values for healthy adult males and females. For both asymmetry magnitude and fat fraction percentage, there was no effect of biologic sex. However, for both variables, a significant effect dependent on muscle was found, indicating that muscle-specific assessment of interlimb imbalances and fat fraction is necessary to properly characterize the musculoskeletal system. These findings align with and expand upon previous research that demonstrated that asymmetry (also quantified by MRI segmentation) varied across quadriceps and hamstring^28^ and that fat fraction percentages varies across muscles of the thigh.^29^

We developed a novel pipeline for utilizing convolutional neural networks to power the segmentation process, which would have been virtually prohibitive if dependent on manual approaches. The CNN approach built upon our prior development and work in the lower extremity^10^, shoulder^12^, and in patients with neuromuscular disease^11^. While the training dataset for the lower extremity was already extensive per prior work, we developed a new training dataset for the trunk and upper extremity muscles. We verified that our approach to segmentation was consistent with prior work by comparing the average volume fraction of each muscle to those seen in literature^6,7^ (**Supplemental Materials Table S10**). All average regional volume fractions were within 5% of literature-reported values. Visually, our segmentation was consistent with high resolution images provided by the Visible Human Project^30^ (our segmentation atlas is shown in **Supplemental Materials Fig. S1**).

As the CNN was re-trained, all muscle labels were reviewed and edited by trained segmentation engineers. We performed both inter- and intra-observer repeatability analyses and found that, on average across, the dice score was 0.94, the volume error was 1.5%, and the fat fraction error was 0.36% fat fraction (**Supplemental Materials Table S1)**. Muscles with the lowest repeatability were the small muscles in the forearms, deep hip and calf, in which small changes to the volume result in large, calculated error. Errors between the final CNN output as compared to a vetted dataset that was not included in training fraction (**Supplemental Materials Table S1)** were similar to the inter- and intra-observer repeatability, illustrating the robustness of the algorithm for future use.

Several limitations of this study should be discussed. First, the study cohort included young-middle-aged individuals, and the amount of ethnic and racial information collected was limited. Therefore, this dataset does not adequately address the impact of age^29^, ethnicity^31^, or race^32^ on muscle or bone volume. Second, precision challenges associated with low volume muscles and/or muscles that are oriented in the in-plane direction (i.e. subclavius, anconeus, piriformis, gemelli, quadratus femoris, obturator internus, obturator externus, and popliteus) lead to increased errors in volume, asymmetry, and fat fraction. To prevent these issues, refined imaging protocols that provide isotropic resolution would improve the precision of these muscles; or, if a refined imaging protocol is not possible, these muscles could be grouped with adjacent or nearby muscle structures to minimize these effects. Third, while muscle volume and fat fraction each have a major influence on muscle force and function, several other factors – such as neuromuscular innervation, pennation angle, optimal fiber length, tendon properties, extracellular matrix properties – also influence muscle force and should be considered within the context of muscle function.

This study presents a comprehensive analysis of 71 muscles bilaterally across the upper body, trunk, and lower body, examining key muscle metrics such as volume, fat fraction, asymmetry, anatomical length, and muscle scaling. We identified novel relationships between muscle size, bone size, and body size, contributing a new reference dataset with broad applications. Muscle volume is a critical indicator of human health^33^ and longevity.^34^ By establishing expected muscle volume based on bone size, height, and mass, deviations from these normative values can be assessed, as has been done previously for lower extremity muscles in both patient populations^7,35,36^ and athletes.^37–39^ This framework holds promise for monitoring changes in muscle over time due to aging, injury, muscle disease, cancer-related cachexia, and a variety of other chronic conditions.^40^

## METHODS

### Dataset

Forty-eight active, healthy subjects (24 females and 24 males with the following subject characteristics (mean +/- standard deviation [range]): age: 29 +/- 8 [19 – 49] years, height: 1.74 +/- 0.10 [1.52 - 2.01] m, body mass: 71.88 +/- 10.15 [48.52 - 92.97] kg) all with no reported history of lower/upper limb injury or surgery within the last 6 months, provided informed consent for this study (**Table 1A**). Subjects were recruited such that the distribution of patients’ heights and weights would reflect those found within the US population, and age was restricted to above 18 and under 50 years old. Subjects were recruited and scanned at one of three distinct imaging sites: Virginia (VA), Ghent University (Belgium), or Wisconsin (WI). All experimental protocols used to collect and process the data were in accordance with relevant guidelines/regulations and were approved by local or central IRBs at University of Virginia, Ghent University, and Advarra.

**Table 1.** Subject demographics and MRI scan parameter details.

| *A. Subject Demographics* |  |  |  |
| --- | --- | --- | --- |
|  | **Males (n = 24)** | **Females (n = 24)** | **All (n = 48)** |
| *Age (years)* | 28 ± 8<br>[19 – 49] | 30 ± 8<br>[19 – 48] | 29 ± 8<br>[19 – 49] |
| *Height (meters)* | 1.81 ± 0.08<br>[1.70 - 2.01] | 1.68 ± 0.07<br>[1.52 - 1.83] | 1.74 ± 0.10<br>[1.52 - 2.01] |
| *Weight (kilograms)* | 77.17 ± 9.04<br>[61.97 - 92.97] | 66.59 ± 8.39<br>[48.52 - 90.70] | 71.88 ± 10.15<br>[48.52 - 92.97] |
| *B. MRI Scan Parameters* |  |  |  |
|  | **VA Site** | **WI Site** | **Ghent Site** |
| *Subject Count* | 8 | 30 | 10 |
| *MRI Scanner* | Siemens Prisma 3T | GE SIGNA Voyager | Siemens MAGNETOM Prisma |
| *TR/TE* | 1.24 & 2.47/4.13 ms | 2.08 & 4.17/6.16 ms | 1.27/4.27 ms |
| *Field of View* | 500 mm x 375 mm | 500 mm x 400 mm | 450 mm x 338 mm |
| *Slice Thickness* | 5 mm | 5 mm | 5 mm |
| *In-Plane Resolution* | 1.3 mm - 2.0 mm | 1.2 mm - 1.7 mm | 1.1 mm – 2 mm |

### Whole body MRI protocol

All MRI scans were acquired using a 3D Axial gradient-echo T1 DIXON sequence (**Table 1B**) in three stages of anatomical coverage: an acquisition of the trunk and lower extremity (LE_EXT), an acquisition of the left side of the upper body (UBL), and an acquisition of the right side of the upper body (UBR). Subjects started the LE_EXT scan with a breath hold to reduce movement artifact of the abdominal muscles, with the scan acquisition covering from the sternum through the mid-pelvis. The LE_EXT scan then was continued with the inherent body coil and no breath hold from the iliac crest through the lateral malleolus. Next, the subjects were shifted off-center on the table, so the right side was up against the bore, the left arm was down at their hip, and the left palm lay flat on the table. Consecutive slabs were then acquired from the subject’s cerebellum to elbow, and then elbow to metacarpals. Images acquired between the elbow and metacarpals had higher in-plane resolution for more detailed visualization of forearm muscle anatomy. The entire left side of the upper body was contained within the field of view. This process was then repeated for the right side (UBR). The spinal column was included in the field of view for both the UBL and UBR acquisitions to ensure continuous left-right coverage. The breath-hold portion of the LE_EXT scan utilized a posterior coil and an anterior torso coil to acquire 20 slices per acquisition slab, while the other sequences utilized the inherent body coil to acquire 40 slices per acquisition slab. Each stage’s MR images were combined to create a single continuous 3D image for the LE_EXT, UBR, and UBL region for both the water and fat phase.

### Muscle and bone segmentation across the full body

In total, 71 muscle structures and 10 bone structures were segmented bilaterally, as well as 3 unilateral bone structures for a total of 165 regions of interest (ROIs) across the three full body MRI regions **(Figure 1)**. Segmentation of the full body was separated into analysis of the LE_EXT, UBR, and UBL regions individually. For the segmentation of the LE_EXT region, we leveraged our previously developed AI-based approach^10^ to segment the boundaries of 87 ROIs, which includes 38 bilateral muscles, 5 bilateral bones and 1 unilateral bone For each AI output, a trained segmentation engineer manually reviewed and refined the segmentations using the segment editor tools in 3D Slicer (v4.11).^41^ Segmentation of the upper body muscles and bones consisted of developing the atlas for segmenting the upper extremity and trunk regions, manually segmenting the muscles, and training a new AI algorithm. The atlas was created through consultation with prior literature^6^ and cross referenced with the Visible Human Data set (from the National Library of Medicine, National Institutes of Health; accessed from https://doi.org/10.37019/e-anatomy/127139). The upper body structures were segmented across UBL and UBR MRI regions of 78 ROIs, comprised of 33 bilateral muscles, 5 bilateral bones and 2 unilateral bones

### AI Algorithm for Segmentation

We utilized an AI-based approach similar to our previously published algorithm^10^ to segment the boundaries of up to 76 muscles (left and right sides) and 14 bones (depending on imaging coverage) of the lower extremity from the Dixon water images. As a brief description, the AI model utilized a modified 3D U-Net structure. Specifically, every level in the encoder contains layers of two blocks of a 3×3×3 convolution layer, a batch normalization (BN) layer, and a rectified linear unit (ReLU) activation layer, followed by a 2×2×2 maxpooling, excluding the bottom-most level. In the decoder, each level consists of layers with a 2×2×2 deconvolution layer, followed by two blocks of a 3×3×3 convolution, a BN, and a ReLU layer. In addition, feature maps from the encoder were concatenated to those of the same resolution in the decoder as the skip connection. The final block of the network contains a 1×1×1 convolution layer to reduce the dimension of the features to match the number of label maps, followed by a pixelwise softmax classifier. The algorithms were implemented based on the framework and training of TensorFlow. During training, weights were initialized randomly from Gaussian distribution and updated with an initial learning rate of 0.01 and the pixelwise dice loss + cross-entropy as the loss function, using the adaptive moment estimation (Adam) optimizer for gradient descent. The initial learning rate was 0.01 and the loss function was pixelwise dice loss + cross entropy + volume error. For a detailed description of our AI model, please see our previous publication.^10^ The AI segmentation algorithm for the UBR and UBL regions was trained to assist in segmentation efforts in a stepwise manner, with AI retraining every 5-10 additionally completed upper body segmentations. All AI segmented outputs were vetted by a trained segmentation engineer. The final upper body algorithm was trained to segment 33 muscles and 7 bones. Initial manual segmentation of UBR and UBL scans from a singular subject required 30 hours. For the final segmentations, the AI assisted segmentation and vetting process required less than three hours.

### Evaluation of Inter-Observer, Intra-Observer, and AI segmentation performance

The consistency and precision of the muscle-by-muscle 3D segmentations were examined in two ways: inter-observer repeatability and intra-observer repeatability. Interobserver is the variability between two different engineers vetting the same scans, and intra-observer is the variability of the same engineer processing the same scan twice. We then analyzed AI performance by comparing the AI output to the standard expert vetted segmentation. To evaluate our dataset, a single full body scan was randomly selected and excluded from all AI trainings to undergo vetting by the same segmentation engineer twice, and by a different engineer once. The output from the AI was then compared to one of the vetted segmentation engineer’s results. This was done for all ROIs on the subject’s scan. Variability was investigated by finding the absolute error in ROI volume, difference in fat fraction, and the dice similarity coefficient (DSC). Inter-observer variability, intra-observer variability and AI performance error (Supplemental Table 2) were low. For inter-/intra-observer repeatability, at least 70 of the 71 muscles had a dice score above 0.8, 67/71 had a volume variability below 5%, and 71/71 had a variability in fat fraction below 2%. For AI performance validation, most muscles that had a volume variability above 5% were small muscles in the upper body. With these muscles having lower overall volumes, even small differences in segmentations can result in a large volume variability. Despite these small muscles having larger volume variability, dice scores remained around 0.8 or greater.

### Quantification of volume, volume fraction, asymmetry, length, fat fraction

The 3D ROIs were used for further analysis. For each ROI, the boundary ROI volume was calculated by summing the total number of pixels labeled for that segmented ROI and multiplied by the pixel’s voxel volume. The boundary volume was utilized in the assessment of asymmetry magnitude, normalized muscle volume, and volume fraction. For the fat fraction calculation, the boundary volume was eroded by one pixel around the entire perimeter in 3D to reduce the impact of bordering subcutaneous or intermuscular fat on the intramuscular fat fraction value. The fat fraction (FF) (%) for each muscle was calculated as: 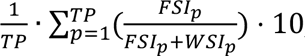, where TP represents the total number of pixels within the muscle, p is an individual pixel, FSI is the pixel’s fat signal intensity and WSI is the pixel’s water signal intensity. Asymmetry between muscles on the left and right side was calculated by subtracting the contralateral muscle volume from the ipsilateral volume and dividing by the sum of left and right muscle volumes. This method results in asymmetry values for right and left muscles with identical magnitudes of opposing signs, with the larger muscle having a positive asymmetry value and the smaller muscle having a negative asymmetry value. The volume percentage of individual muscles relative to total muscle volume across the full body was determined for subjects that had the requisite muscle coverage (Males: n=13; Females: n=20) to investigate differences in how muscle volumes are distributed based on biological sex.

Muscle anatomical length was found as described in our previous work using axial centroids of the 3D segmentation^5^. For each muscle segmentation, the centroid of each muscle label was computed on each axial slice. The 3D Euclidean distance between adjacent-slice centroids was calculated and all inter-slice distances were summed to yield muscle length. By defining muscle length using centroids, lengths represent the anatomical line-of-action, incorporating the muscle’s curved shape and complex wrapping. Bone length measures were found by determining the Euclidean distance between two anatomical landmarks located on 3D bone segmentations. The following bone lengths were found: humerus length (superior aspect of the humeral head to inferior aspect of the trochlea), ulna length (superior aspect of the olecranon process to the inferior aspect of the styloid process), vertebra length (top of first vertebra to bottom of last vertebra), pelvic height (superior aspect of the ilium to the inferior aspect of the pubis arch), femur length (superior aspect of femoral head to inferior aspect of medial condyle), and tibia length (tibial plateau to medial malleolus).

For analysis purposes, muscle scaling relationships, size, asymmetry, and fat fraction were investigated at the individual and group levels. In total, 19 functional groups and 7 joint groups were defined; each of the 71 unique muscles was included in only one functional/joint group (thus biarticular muscles or muscles that cross joints with multiple degrees of freedom were only included in one functional/joint group). Separate from the functional groups, the total upper extremity, lower extremity, and full body muscle groupings were also investigated. The lower body consisted of all muscles below the iliac crest (including the psoas major) and the upper body consisted of all muscles above the iliac crest.

### Volume, length, distribution, asymmetry, and fat fraction across muscles

Two-way mixed repeated ANOVAs were used to assess differences in asymmetry magnitude, intramuscular fat fraction percentage, normalized volume, volume fraction, and difference between actual and predicted volume using varying linear regression models between biological sexes and muscle structures. Biological sex was the between-subject effect and muscle was the within-subject effect. The within-subject effect of muscle was run on individual muscles and functional muscle groupings for all dependent variables except for volume fraction and difference between actual and predicted volumes, which were run only on individual muscles.. A log10 transform was applied to asymmetry, fat infiltration, normalized muscle volume, and volume fraction datasets to adhere to the assumption of normality. All significant main effects were reported, and any interaction effects were investigated using post-hoc pairwise t-test analyses. A statistical significance level of α = 0.05 was used.

### Cluster analysis of muscle volume distribution

Hierarchical clustering analysis was used to examine the patterns of muscle distribution across the population, similar to that described in Knaus et al.^38^ For each muscle, the right and left muscle volumes were summed and represented as a fraction of the total muscle volume; a z-score was then calculated for each muscle by comparing with the average volume fraction of that muscle (across all individuals) and representing that difference in units of standard deviation across the individuals. The result was a 71-by-37 matrix of Z-scores: 71 muscles and 37 subjects (only subjects with full muscle coverage were included).

The clustering analysis then was performed to understand the similarity in muscle distribution patterns. Each muscle is described as a vector, defined by row values, that exists in multi-dimensional space in which each dimension is defined by one athlete limb. Therefore, “subject space” has as many dimensions as the number of subjects included (n=37), and as many vectors as there are muscles (n=71) exist in this space. Muscle vectors were clustered according to their Euclidean distance in “subject space”, meaning vectors with the most similar orientation and magnitude cluster. Vectors were linked in pairs to build a hierarchy; all vectors were compared and the two closest were clustered together. The average of this pair defined a new vector and was then compared with all remaining vectors to identify the next closest pair, repeating the process until all the original vectors were linked. These linkages are illustrated by a hierarchical dendrogram, or tree diagram. The same clustering process was also applied to the subjects (columns). Now columns define vectors in “muscle space” with as many dimensions as there are muscles (n=71) and in this space exists as many vectors as there are subjects (n=37). Hierarchical clustering of muscles was determined from the Euclidian distance of these vectors in “muscle space.” The result is rows and columns of the original data matrix rearranged based on clustering, depicted with their dendrograms.

### Muscle volume and length scaling

Scaling relationships were examined by performing linear regression analyses between muscle volumes and body/bone dimensions. Scaling analyses were performed at the individual muscle, muscle group, upper vs. lower body, and total body levels. Body dimensions examined included: body mass, height, body mass index (BMI) and the product of body mass and height.^5^ Bone dimensions examined included volume and length on the bone in closest proximity to each muscle (**Table 2**). The average of left and right muscle volumes was correlated via a linear regression to total muscle volume, body mass, height, BMI, mass*height, and the associated bone volume. All left and right muscle lengths were correlated via a linear regression to their respective bone length. Similar analyses were performed for total muscle volume/total bone volume, total upper body muscle volume/upper bone volume, and total lower body muscle volume/lower body bone volume.

To statistically compare the scaling relationships, we found the error between the actual to predicted volume for each muscle and each method (total body muscle volume, corresponding bone volume, height, mass, and height*mass). A two-way repeated ANOVA was used with muscle and scaling methods utilizing the prediction error as the dependent variable. For analyzing mass*height scaling, the effect of sex was investigated with a two-way mixed repeated ANOVA, where biological sex was the between-subject effect and muscle was the within-subject effect. All significant main effects and/or interaction effects were investigated using post-hoc pairwise t-test analyses. Lastly, to statistically compare the scaling relationship of corresponding bone length on muscle length and the effect of sex, we found the error between the actual to predicted muscle length for each muscle. A two-way mixed repeated ANOVA was used with muscle as the within subject effects and sex as the between subject effect utilizing the prediction error as the dependent variable. A statistical significance level of α = 0.05 was used. The *pingouin* package^42^ (v0.5.4) for python (v3.8.10) was used for all statistical analyses.

### Analysis of deviations from scaling relationships

To explore the deviations from expected muscle size that may vary across parameters, we quantified the errors between measured and predicted muscle volume and expressed these errors as a z-score from the average value. This analysis was performed for the muscle volume vs. height relationship, the muscle volume vs. associated bone volume relationship, and the bone volume vs. height relationship. The analysis was repeated at the total muscle/total bone and individual muscle/associated bone levels. The body parameter of height was chosen to avoid the confounding effect of muscle volume on body mass since body mass will naturally increase as muscle volume increases.

## Author Contributions Statement

LR, MP, OD, and SSB contributed to study design, analysis, data interpretation, and paper writing. TB, XF, MC, SSB, and LR contributed to MRI protocol initialization and data collection. LR, MP, OD, MC, and SDB assisted in study design, data interpretation, and revision of the paper. MP, OD, MC, VP, JM, AC, LH, RH, MR, and EC assisted in image segmentation and LR, XF in assistive automation. All authors have read and approved the final submitted manuscript.

## Additional Information

Financial competing interests: LR, MP, OD, VP, JM, AC, LH, EC, MC, XF, SDB, TB, RH, MR, and SSB are compensated by Springbok Analytics either as an employee or consultant. LR, MP, MC, AC, SDB, JM, TB, and OD have stock options to the company. XF and SSB own stock in the company.

## Data Sharing

Data generated or analyzed during the study are available from the corresponding author by request.

