## Supplemental information for "Big Bones Mean Big Muscles: AI Quantifies 71 Individual Muscles Across the Body, Revealing Widespread Links Between Muscle, Bone, and Size"

### <u>Supplemental Material</u>

Supplemental material contains the following:


- **Segmentation Repeatability Validation** (Page 2): Detailed results, statistics, and table for the segmentation repeatability validation.
- **Muscle Volume Scaling** (Page 5): Detailed results and statistics for the two-way repeated ANOVA to investigate muscle volume scaling methods as the within subject effects utilizing the prediction error as the dependent variable. Additionally, within the product method, the effect of sex was investigated with a two-way mixed repeated ANOVA.
- **Muscle Length Scaling** (Page 6): Detailed results and statistics for the two-way mixed repeated ANOVA to investigate the effect of sex and muscle on the prediction error of muscle length using bone length scaling as the dependent variable.
- **Analysis of Metric Variation Across Muscles and Biologic Sex** (Page 7): Detailed results and statistics for the two-way mixed repeated ANOVA to investigate the effect of sex and muscle on asymmetry, fat infiltration, normalized muscle volume, muscle volume fraction, and the difference between actual and predicted volume.
- **Segmentation Atlas (**Page 15): Upper Body and Lower Extremity segmentation atlas.
- **Comparison to Literature** (Page 17): Detailed methods and table for average muscle volume fractions compared to literature for Upper Body and Lower Extremity muscles.

### Segmentation Repeatability Validation

Overall, all ROIs displayed low interobserver and intraobserver variability, as well as AI validation error being low (**Table S1**). Interobserver and interobserver repeatability measures were similar. When the ROI results were averaged for the right and left side, at least 70 of the 71 muscles had a dice score above 0.8, 67/71 had a volume error below 5%, and 70/71 had a difference in fat fraction below 2% for both inter and intraobserver repeatability. Further, at least 55 of the 71 muscles had a dice score above 0.9, 63/71 had a volume error below 3%, and 65/71 had a difference in fat fraction below 1% for both interobserver and intraobserver repeatability. The ROIs with the lowest repeatability error were usually large muscles (latissimus dorsi, triceps brachii, erector spinae, gluteus maximus, and vastus lateralis all had a dice > .98, volume error below 1%, and fat fraction difference below 1%), while smaller muscles in which very minute changes to the segmentation caused a higher relative change had the largest repeatability errors (flexor carpi radialis and palmaris longus, flexor digitorum superficialis, flexor digitorum profundus and flexor hallucis longus had dice scores ~ 0.8 and volume errors above 5%). There seemed to be no trend between the upper and lower body segmentation repeatability. This seemed to be repeated by the AI validation results, in which averaged ROIs for the right and left side resulted in: at least 70 of the 71 muscles with a dice score above 0.8, 63/71 with a volume error below 5%, and 71/71 with a difference in fat fraction below 2%. Further, at least 55 of the 71 muscles had a dice score above 0.9, 57/71 had a volume error below 3%, and 61/71 had a difference in fat fraction below 1%.

**Table S1**: The Dice Score (DSC), volume error (%) and fat fraction difference (%) for the interobserver and intraobserver repeatability comparison for each muscle/bone.

| Muscle Name | Dice Score (DSC) | | | Volume Error (%) | | | Fat Fraction Difference (%) | | |
|---|---|---|---|---|---|---|---|---|---|
| | Inter | Intra | AI | Inter | Intra | AI | Inter | Intra | AI |
| levator scapulae | 0.905 ± 0.030 | 0.924 ± 0.036 | 0.915 ± 0.045 | 2.60 ± 0.115 | 3.67 ± 3.590 | 5.07 ± 4.649 | 0.74 ± 0.343 | 0.29 ± 0.254 | 0.48 ± 0.339 |
| supraspinatus | 0.950 ± 0.018 | 0.944 ± 0.004 | 0.945 ± 0.029 | 2.46 ± 1.797 | 2.64 ± 0.739 | 3.05 ± 2.919 | 0.08 ± 0.039 | 0.20 ± 0.085 | 0.15 ± 0.086 |
| trapezius | 0.969 ± 0.005 | 0.965 ± 0.002 | 0.965 ± 0.009 | 0.78 ± 0.579 | 1.31 ± 0.259 | 1.30 ± 1.132 | 0.23 ± 0.011 | 0.57 ± 0.104 | 0.51 ± 0.377 |
| rhomboid | 0.909 ± 0.001 | 0.937 ± 0.014 | 0.908 ± 0.000 | 7.22 ± 0.326 | 3.56 ± 1.802 | 7.44 ± 0.050 | 0.23 ± 0.064 | 0.18 ± 0.156 | 0.30 ± 0.018 |
| serratus anterior | 0.957 ± 0.016 | 0.978 ± 0.007 | 0.980 ± 0.007 | 1.49 ± 0.598 | 0.36 ± 0.217 | 1.09 ± 0.539 | 0.57 ± 0.307 | 0.02 ± 0.015 | 0.09 ± 0.001 |
| latissimus dorsi | 0.988 ± 0.002 | 0.990 ± 0.002 | 0.992 ± 0.002 | 0.53 ± 0.409 | 0.37 ± 0.161 | 0.43 ± 0.237 | 0.23 ± 0.176 | 0.09 ± 0.033 | 0.17 ± 0.090 |
| subscapularis | 0.977 ± 0.005 | 0.977 ± 0.002 | 0.978 ± 0.003 | 0.56 ± 0.022 | 0.66 ± 0.212 | 1.08 ± 0.350 | 0.15 ± 0.112 | 0.19 ± 0.059 | 0.23 ± 0.209 |
| teres major | 0.942 ± 0.001 | 0.941 ± 0.001 | 0.942 ± 0.001 | 0.38 ± 0.346 | 0.32 ± 0.113 | 0.38 ± 0.346 | 0.02 ± 0.001 | 0.03 ± 0.012 | 0.02 ± 0.001 |
| infraspinatus and teres minor | 0.966 ± 0.021 | 0.982 ± 0.006 | 0.980 ± 0.007 | 1.38 ± 1.346 | 0.20 ± 0.067 | 0.39 ± 0.355 | 0.17 ± 0.108 | 0.04 ± 0.025 | 0.10 ± 0.040 |
| subclavius | 0.833 ± 0.010 | 0.831 ± 0.008 | 0.847 ± 0.024 | 1.61 ± 1.607 | 1.61 ± 1.609 | 0.00 ± 0.000 | 0.18 ± 0.177 | 0.01 ± 0.006 | 0.00 ± 0.000 |
| pectoralis minor | 0.965 ± 0.001 | 0.964 ± 0.000 | 0.965 ± 0.001 | 0.73 ± 0.295 | 0.60 ± 0.190 | 0.72 ± 0.291 | 0.12 ± 0.033 | 0.11 ± 0.073 | 0.12 ± 0.034 |
| pectoralis major | 0.974 ± 0.015 | 0.982 ± 0.011 | 0.975 ± 0.019 | 0.28 ± 0.087 | 0.73 ± 0.507 | 0.90 ± 0.710 | 0.18 ± 0.146 | 0.30 ± 0.118 | 0.02 ± 0.016 |
| deltoid | 0.987 ± 0.009 | 0.992 ± 0.005 | 0.988 ± 0.008 | 0.89 ± 0.877 | 0.27 ± 0.261 | 0.80 ± 0.789 | 0.03 ± 0.027 | 0.00 ± 0.002 | 0.02 ± 0.019 |
| triceps brachii | 0.993 ± 0.001 | 0.992 ± 0.002 | 0.993 ± 0.001 | 0.26 ± 0.004 | 0.28 ± 0.197 | 0.28 ± 0.015 | 0.01 ± 0.001 | 0.23 ± 0.218 | 0.00 ± 0.002 |
| coracobrachialis | 0.960 ± 0.000 | 0.960 ± 0.000 | 0.960 ± 0.000 | 0.00 ± 0.000 | 0.00 ± 0.000 | 0.00 ± 0.000 | 0.00 ± 0.000 | 0.00 ± 0.000 | 0.00 ± 0.000 |
| biceps brachii | 0.987 ± 0.003 | 0.987 ± 0.003 | 0.987 ± 0.003 | 0.32 ± 0.317 | 0.31 ± 0.308 | 0.32 ± 0.321 | 0.25 ± 0.247 | 0.18 ± 0.179 | 0.25 ± 0.251 |

| | | | | | | | | | |
|---|---|---|---|---|---|---|---|---|---|
| **brachialis** | 0.972 ± 0.004 | 0.978 ± 0.006 | 0.975 ± 0.008 | 1.11 ± 0.036 | 0.32 ± 0.201 | 0.54 ± 0.198 | 0.25 ± 0.085 | 0.04 ± 0.011 | 0.17 ± 0.046 |
| **brachioradialis** | 0.944 ± 0.011 | 0.955 ± 0.003 | 0.945 ± 0.012 | 2.40 ± 1.383 | 1.44 ± 0.137 | 3.10 ± 0.681 | 0.14 ± 0.063 | 0.25 ± 0.248 | 0.23 ± 0.022 |
| **extensor carpi radialis** | 0.945 ± 0.001 | 0.946 ± 0.011 | 0.947 ± 0.001 | 1.96 ± 0.512 | 1.60 ± 0.371 | 1.70 ± 0.607 | 1.49 ± 0.255 | 0.78 ± 0.531 | 1.45 ± 0.287 |
| **anconeus** | 0.798 ± 0.014 | 0.805 ± 0.022 | 0.798 ± 0.014 | 4.00 ± 0.380 | 0.63 ± 0.361 | 4.00 ± 0.380 | 0.11 ± 0.032 | 0.02 ± 0.009 | 0.11 ± 0.032 |
| **digit extensors** | 0.860 ± 0.045 | 0.865 ± 0.036 | 0.848 ± 0.044 | 2.05 ± 1.227 | 3.35 ± 2.335 | 4.60 ± 1.867 | 2.15 ± 0.969 | 1.14 ± 0.044 | 1.15 ± 0.576 |
| **extensor carpi ulnaris** | 0.843 ± 0.002 | 0.859 ± 0.004 | 0.839 ± 0.003 | 2.25 ± 0.192 | 2.80 ± 0.906 | 6.39 ± 0.868 | 0.55 ± 0.460 | 0.54 ± 0.012 | 1.12 ± 0.412 |
| **deep forearm extensors** | 0.895 ± 0.026 | 0.901 ± 0.021 | 0.886 ± 0.031 | 1.59 ± 0.388 | 0.43 ± 0.244 | 1.49 ± 1.302 | 0.78 ± 0.386 | 0.77 ± 0.336 | 1.37 ± 0.763 |
| **supinator** | 0.887 ± 0.004 | 0.880 ± 0.014 | 0.890 ± 0.001 | 1.30 ± 0.305 | 1.06 ± 0.950 | 1.13 ± 1.022 | 0.23 ± 0.206 | 0.05 ± 0.036 | 0.20 ± 0.183 |
| **pronator teres** | 0.936 ± 0.001 | 0.944 ± 0.003 | 0.938 ± 0.005 | 0.81 ± 0.474 | 0.22 ± 0.143 | 0.95 ± 0.150 | 0.74 ± 0.319 | 0.19 ± 0.174 | 0.52 ± 0.480 |
| **flexor carpi radialis & palmaris longus** | 0.940 ± 0.004 | 0.873 ± 0.067 | 0.873 ± 0.070 | 1.68 ± 0.619 | 6.70 ± 5.125 | 6.24 ± 5.865 | 0.63 ± 0.536 | 1.67 ± 0.507 | 1.53 ± 0.578 |
| **flexor digitorum superficialis** | 0.958 ± 0.006 | 0.821 ± 0.141 | 0.818 ± 0.140 | 1.60 ± 0.604 | 15.24 ± 15.07 | 16.16 ± 14.49 | 0.33 ± 0.181 | 0.39 ± 0.312 | 0.57 ± 0.089 |
| **flexor carpi ulnaris** | 0.931 ± 0.002 | 0.929 ± 0.012 | 0.922 ± 0.012 | 2.42 ± 0.860 | 1.52 ± 0.249 | 3.82 ± 0.498 | 0.32 ± 0.191 | 0.70 ± 0.655 | 0.40 ± 0.232 |
| **flexor digitorum profundus** | 0.966 ± 0.005 | 0.969 ± 0.005 | 0.971 ± 0.002 | 0.60 ± 0.373 | 0.40 ± 0.151 | 0.74 ± 0.119 | 0.06 ± 0.055 | 0.02 ± 0.013 | 0.08 ± 0.024 |
| **flexor pollicis longus** | 0.896 ± 0.020 | 0.840 ± 0.061 | 0.841 ± 0.062 | 1.77 ± 0.484 | 8.37 ± 6.952 | 7.82 ± 6.964 | 0.28 ± 0.265 | 0.51 ± 0.491 | 0.51 ± 0.475 |
| **pronator quadratus** | 0.830 ± 0.001 | 0.852 ± 0.003 | 0.833 ± 0.002 | 4.81 ± 0.328 | 1.60 ± 1.046 | 4.57 ± 0.041 | 0.49 ± 0.159 | 0.18 ± 0.168 | 0.41 ± 0.198 |
| **rectus abdominis** | 0.971 ± 0.002 | 0.961 ± 0.008 | 0.937 ± 0.005 | 0.67 ± 0.489 | 1.23 ± 0.768 | 3.83 ± 0.239 | 0.17 ± 0.070 | 0.18 ± 0.173 | 0.16 ± 0.128 |
| **external oblique** | 0.926 ± 0.002 | 0.931 ± 0.004 | 0.884 ± 0.005 | 3.68 ± 0.179 | 2.80 ± 0.262 | 8.09 ± 0.581 | 0.43 ± 0.156 | 0.28 ± 0.223 | 0.25 ± 0.101 |
| **erector spinae** | 0.987 ± 0.002 | 0.993 ± 0.000 | 0.992 ± 0.002 | 0.96 ± 0.186 | 0.33 ± 0.055 | 0.51 ± 0.117 | 0.06 ± 0.004 | 0.01 ± 0.001 | 0.01 ± 0.002 |
| **multifidus** | 0.979 ± 0.006 | 0.985 ± 0.005 | 0.973 ± 0.016 | 0.83 ± 0.175 | 0.43 ± 0.353 | 2.06 ± 1.595 | 0.12 ± 0.091 | 0.13 ± 0.117 | 0.15 ± 0.099 |
| **quadratus lumborum** | 0.899 ± 0.024 | 0.915 ± 0.014 | 0.910 ± 0.000 | 0.95 ± 0.725 | 1.52 ± 0.226 | 2.45 ± 2.074 | 0.31 ± 0.104 | 0.10 ± 0.032 | 1.05 ± 0.642 |
| **psoas major** | 0.968 ± 0.002 | 0.967 ± 0.002 | 0.979 ± 0.000 | 0.64 ± 0.272 | 0.76 ± 0.159 | 0.15 ± 0.006 | 0.25 ± 0.234 | 0.27 ± 0.217 | 0.04 ± 0.038 |
| **iliacus** | 0.959 ± 0.002 | 0.959 ± 0.003 | 0.969 ± 0.001 | 0.90 ± 0.332 | 0.73 ± 0.223 | 0.21 ± 0.115 | 0.31 ± 0.026 | 0.28 ± 0.055 | 0.16 ± 0.089 |
| **gluteus medius** | 0.979 ± 0.001 | 0.978 ± 0.001 | 0.983 ± 0.000 | 0.65 ± 0.063 | 0.56 ± 0.037 | 0.18 ± 0.072 | 0.21 ± 0.040 | 0.20 ± 0.079 | 0.09 ± 0.064 |
| **gluteus maximus** | 0.988 ± 0.000 | 0.988 ± 0.000 | 0.992 ± 0.000 | 0.09 ± 0.040 | 0.08 ± 0.033 | 0.19 ± 0.016 | 0.15 ± 0.059 | 0.14 ± 0.052 | 0.13 ± 0.009 |
| **gluteus minimus** | 0.941 ± 0.001 | 0.942 ± 0.000 | 0.951 ± 0.001 | 1.30 ± 0.412 | 1.15 ± 0.228 | 0.73 ± 0.151 | 0.09 ± 0.054 | 0.10 ± 0.065 | 0.12 ± 0.023 |
| **piriformis** | 0.896 ± 0.011 | 0.889 ± 0.008 | 0.920 ± 0.007 | 2.91 ± 0.218 | 3.63 ± 0.416 | 1.57 ± 0.085 | 0.95 ± 0.343 | 1.19 ± 0.352 | 0.48 ± 0.052 |
| **gemelli** | 0.834 ± 0.019 | 0.836 ± 0.019 | 0.852 ± 0.013 | 2.52 ± 0.210 | 1.09 ± 0.980 | 1.32 ± 0.555 | 1.17 ± 0.133 | 0.69 ± 0.409 | 0.30 ± 0.274 |
| **quadratus femoris** | 0.888 ± 0.005 | 0.885 ± 0.003 | 0.899 ± 0.007 | 0.55 ± 0.292 | 0.46 ± 0.282 | 0.15 ± 0.066 | 0.31 ± 0.212 | 0.26 ± 0.172 | 0.22 ± 0.023 |
| **obturator internus** | 0.843 ± 0.004 | 0.844 ± 0.004 | 0.869 ± 0.010 | 1.60 ± 1.293 | 0.94 ± 0.055 | 0.58 ± 0.171 | 0.38 ± 0.247 | 0.31 ± 0.176 | 0.31 ± 0.106 |
| **obturator externus** | 0.906 ± 0.006 | 0.905 ± 0.006 | 0.913 ± 0.004 | 1.33 ± 0.643 | 1.56 ± 0.674 | 2.79 ± 0.779 | 0.16 ± 0.123 | 0.13 ± 0.115 | 0.15 ± 0.019 |
| **pectineus** | 0.934 ± 0.001 | 0.934 ± 0.001 | 0.947 ± 0.001 | 0.96 ± 0.872 | 0.96 ± 0.872 | 0.54 ± 0.010 | 0.32 ± 0.063 | 0.32 ± 0.063 | 0.19 ± 0.083 |
| **tensor fasciae latae** | 0.923 ± 0.005 | 0.935 ± 0.007 | 0.931 ± 0.005 | 1.88 ± 1.013 | 0.74 ± 0.592 | 1.38 ± 0.470 | 0.99 ± 0.931 | 0.05 ± 0.041 | 1.05 ± 0.865 |
| **rectus femoris** | 0.980 ± 0.002 | 0.980 ± 0.002 | 0.984 ± 0.001 | 0.12 ± 0.086 | 0.14 ± 0.073 | 0.11 ± 0.057 | 0.04 ± 0.025 | 0.04 ± 0.027 | 0.00 ± 0.002 |
| **vastus lateralis** | 0.989 ± 0.001 | 0.989 ± 0.001 | 0.992 ± 0.000 | 0.14 ± 0.086 | 0.18 ± 0.025 | 0.08 ± 0.006 | 0.13 ± 0.054 | 0.12 ± 0.048 | 0.05 ± 0.021 |
| **vastus intermedius** | 0.975 ± 0.003 | 0.975 ± 0.003 | 0.980 ± 0.001 | 0.70 ± 0.262 | 0.70 ± 0.262 | 0.20 ± 0.038 | 0.05 ± 0.005 | 0.05 ± 0.005 | 0.01 ± 0.003 |
| **vastus medialis** | 0.987 ± 0.001 | 0.987 ± 0.001 | 0.989 ± 0.001 | 0.11 ± 0.065 | 0.11 ± 0.061 | 0.05 ± 0.016 | 0.10 ± 0.053 | 0.10 ± 0.050 | 0.03 ± 0.027 |

| | | | | | | | | | |
|---|---|---|---|---|---|---|---|---|---|
| **sartorius** | 0.957 ± 0.001 | 0.961 ± 0.003 | 0.964 ± 0.002 | 1.71 ± 0.063 | 1.23 ± 0.368 | 0.36 ± 0.344 | 0.46 ± 0.179 | 0.21 ± 0.117 | 0.32 ± 0.221 |
| **adductor brevis** | 0.934 ± 0.002 | 0.933 ± 0.002 | 0.948 ± 0.002 | 0.44 ± 0.393 | 0.33 ± 0.286 | 0.61 ± 0.152 | 0.14 ± 0.044 | 0.13 ± 0.026 | 0.02 ± 0.017 |
| **adductor magnus** | 0.983 ± 0.000 | 0.983 ± 0.000 | 0.987 ± 0.000 | 0.25 ± 0.011 | 0.25 ± 0.011 | 0.08 ± 0.048 | 0.04 ± 0.008 | 0.05 ± 0.011 | 0.08 ± 0.049 |
| **adductor longus** | 0.965 ± 0.001 | 0.965 ± 0.001 | 0.968 ± 0.000 | 0.62 ± 0.320 | 0.62 ± 0.320 | 0.92 ± 0.153 | 0.06 ± 0.034 | 0.06 ± 0.034 | 0.07 ± 0.001 |
| **gracilis** | 0.939 ± 0.004 | 0.944 ± 0.003 | 0.948 ± 0.004 | 1.07 ± 0.750 | 0.75 ± 0.393 | 0.39 ± 0.144 | 0.97 ± 0.027 | 0.40 ± 0.084 | 0.65 ± 0.064 |
| **semitendinosus** | 0.976 ± 0.003 | 0.976 ± 0.003 | 0.979 ± 0.001 | 0.25 ± 0.086 | 0.28 ± 0.018 | 0.18 ± 0.128 | 0.13 ± 0.009 | 0.12 ± 0.087 | 0.02 ± 0.004 |
| **semimembranosus** | 0.974 ± 0.002 | 0.974 ± 0.002 | 0.979 ± 0.002 | 0.46 ± 0.125 | 0.42 ± 0.168 | 0.18 ± 0.025 | 0.10 ± 0.071 | 0.09 ± 0.063 | 0.10 ± 0.068 |
| **biceps femoris: long head** | 0.976 ± 0.000 | 0.977 ± 0.000 | 0.978 ± 0.000 | 0.39 ± 0.099 | 0.28 ± 0.059 | 0.15 ± 0.150 | 0.21 ± 0.048 | 0.10 ± 0.078 | 0.08 ± 0.032 |
| **biceps femoris: short head** | 0.945 ± 0.005 | 0.947 ± 0.005 | 0.953 ± 0.001 | 0.63 ± 0.592 | 0.65 ± 0.388 | 0.49 ± 0.282 | 0.37 ± 0.157 | 0.22 ± 0.180 | 0.06 ± 0.014 |
| **popliteus** | 0.839 ± 0.033 | 0.863 ± 0.018 | 0.856 ± 0.026 | 5.50 ± 3.274 | 2.34 ± 1.214 | 2.83 ± 2.243 | 0.36 ± 0.188 | 0.43 ± 0.139 | 0.29 ± 0.224 |
| **gastrocnemius: medial head** | 0.953 ± 0.005 | 0.966 ± 0.002 | 0.960 ± 0.005 | 2.15 ± 0.877 | 0.51 ± 0.014 | 1.74 ± 0.789 | 1.35 ± 0.852 | 0.38 ± 0.059 | 1.11 ± 0.824 |
| **gastrocnemius: lateral head** | 0.955 ± 0.001 | 0.960 ± 0.000 | 0.962 ± 0.004 | 0.54 ± 0.348 | 0.30 ± 0.049 | 0.75 ± 0.282 | 0.21 ± 0.103 | 0.15 ± 0.143 | 0.13 ± 0.121 |
| **soleus** | 0.970 ± 0.003 | 0.972 ± 0.002 | 0.978 ± 0.001 | 0.73 ± 0.268 | 0.28 ± 0.157 | 0.03 ± 0.020 | 0.19 ± 0.035 | 0.05 ± 0.007 | 0.14 ± 0.045 |
| **tibialis anterior** | 0.951 ± 0.002 | 0.958 ± 0.003 | 0.960 ± 0.001 | 0.97 ± 0.034 | 0.30 ± 0.047 | 0.94 ± 0.398 | 0.12 ± 0.057 | 0.07 ± 0.067 | 0.19 ± 0.021 |
| **phalangeal extensors** | 0.932 ± 0.005 | 0.937 ± 0.004 | 0.945 ± 0.001 | 2.01 ± 0.928 | 1.30 ± 0.885 | 1.16 ± 0.538 | 0.27 ± 0.106 | 0.16 ± 0.011 | 0.21 ± 0.095 |
| **fibulari** | 0.945 ± 0.014 | 0.952 ± 0.007 | 0.960 ± 0.005 | 2.24 ± 2.093 | 1.51 ± 1.379 | 0.87 ± 0.761 | 0.23 ± 0.008 | 0.19 ± 0.044 | 0.05 ± 0.025 |
| **tibialis posterior** | 0.949 ± 0.002 | 0.952 ± 0.002 | 0.958 ± 0.001 | 1.12 ± 0.407 | 1.53 ± 0.390 | 0.18 ± 0.019 | 0.07 ± 0.020 | 0.07 ± 0.017 | 0.18 ± 0.018 |
| **flexor digitorum longus** | 0.838 ± 0.003 | 0.857 ± 0.001 | 0.849 ± 0.014 | 3.28 ± 2.454 | 2.36 ± 1.919 | 5.08 ± 0.826 | 1.90 ± 0.243 | 0.22 ± 0.045 | 1.85 ± 0.385 |
| **flexor hallucis longus** | 0.897 ± 0.001 | 0.898 ± 0.001 | 0.935 ± 0.002 | 6.10 ± 0.051 | 6.11 ± 0.057 | 0.59 ± 0.174 | 0.19 ± 0.067 | 0.19 ± 0.062 | 0.10 ± 0.061 |
| **clavicle** | 0.914 ± 0.037 | 0.939 ± 0.013 | 0.917 ± 0.035 | 1.54 ± 1.542 | 1.27 ± 1.270 | 1.74 ± 1.737 | - | - | - |
| **sternum** | 0.968 ± 0.001 | 0.969 ± 0.001 | 0.969 ± 0.001 | 0.01 ± 0.013 | 0.04 ± 0.039 | 0.07 ± 0.070 | - | - | - |
| **ribs** | 0.922 ± 0.014 | 0.939 ± 0.023 | 0.934 ± 0.024 | 2.16 ± 1.684 | 0.48 ± 0.381 | 5.71 ± 2.305 | - | - | - |
| **humerus** | 0.991 ± 0.000 | 0.991 ± 0.000 | 0.991 ± 0.000 | 0.00 ± 0.000 | 0.00 ± 0.000 | 0.00 ± 0.000 | - | - | - |
| **ulna** | 0.968 ± 0.000 | 0.968 ± 0.000 | 0.968 ± 0.000 | 0.00 ± 0.000 | 0.00 ± 0.000 | 0.00 ± 0.000 | - | - | - |
| **radius** | 0.967 ± 0.001 | 0.969 ± 0.000 | 0.969 ± 0.000 | 0.13 ± 0.132 | 0.03 ± 0.029 | 0.00 ± 0.000 | - | - | - |
| **vertebrae** | 0.831 ± 0.031 | 0.844 ± 0.015 | 0.809 ± 0.013 | 4.92 ± 1.941 | 0.92 ± 0.495 | 18.94 ± 1.34 | - | - | - |
| **sacrum** | 0.947 | 0.947 | 0.950 | 0.26 | 0.33 | 1.80 | - | - | - |
| **pelvis** | 0.938 ± 0.004 | 0.931 ± 0.005 | 0.960 ± 0.003 | 3.01 ± 0.635 | 1.08 ± 0.009 | 1.40 ± 0.091 | - | - | - |
| **femur** | 0.980 ± 0.001 | 0.980 ± 0.000 | 0.986 ± 0.000 | 0.53 ± 0.011 | 0.44 ± 0.071 | 0.35 ± 0.048 | - | - | - |
| **patella** | 0.837 ± 0.006 | 0.837 ± 0.006 | 0.843 ± 0.011 | 0.31 ± 0.108 | 0.12 ± 0.084 | 0.26 ± 0.142 | - | - | - |
| **tibia** | 0.966 ± 0.003 | 0.966 ± 0.001 | 0.968 ± 0.013 | 0.92 ± 0.083 | 0.27 ± 0.180 | 1.51 ± 1.251 | - | - | - |
| **fibula** | 0.811 ± 0.044 | 0.809 ± 0.045 | 0.888 ± 0.014 | 3.53 ± 2.113 | 1.67 ± 0.782 | 0.53 ± 0.441 | - | - | - |
| **Average Across All Muscles** | **0.937 ± 0.048** | **0.937 ± 0.050** | **0.937 ± 0.051** | **1.49 ± 1.42** | **3.67 ± 2.252** | **1.94 ± 3.128** | **0.36 ± 0.43** | **0.25 ± 0.30** | **0.32 ± 0.413** |

### Muscle Volume Scaling

*Comparison between Muscle Volume Scaling Methods*

Despite slight variation is correlation coefficient strength between muscles and scaling method, when comparing the error in predicted muscle volume utilizing the three differing normalization method's linear regression across muscles, there were no statistically significant main effect of muscle ($F(70, 2520) = 1.256$, $p_{GG} = 0.322$, $\eta_2 = 0.031$, $\epsilon = 0.122$) or normalization method ($F(2, 72) = 1.044$, $p = 0.357$, $\eta_2 = 0.0282$, $\epsilon = 0.977$), but there was a significant interaction effect ($F(140, 5040) = 10.736$, $p_{GG} < 0.001$, $\eta_2 = 0.230$, $\epsilon = 0.047$). However, post-hoc analysis revealed all significant interaction effects were from comparing muscles within a method. See **Table S2** below for the post-hoc results.

**Table S2**: Post-hoc analysis results for the interaction effect. All comparisons had a degree of freedom of 36, were paired, parametric, with a two-sided alternative hypothesis and used a sidak adjustment. The contrast was Normalization Method*Muscle for all comparisons.

| Normalization Method | A | B | T | p-corr | cohen |
|---|---|---|---|---|---|
| Bone Volume | extensor carpi ulnaris | supinator | -6.156 | 0.003 | -1.176 |
| Product | extensor carpi ulnaris | supinator | -5.992 | 0.005 | -1.078 |
| Bone Volume | extensor carpi ulnaris | flexor carpi radialis &palmaris longus | -5.914 | 0.007 | -0.954 |
| Bone Volume | infraspinatus and teres minor | supinator | -5.718 | 0.012 | -0.914 |
| Product | quadratus lumborum | supinator | -5.480 | 0.025 | -0.862 |
| Product | extensor carpi ulnaris | flexor carpi radialis and palmaris longus | -5.451 | 0.028 | -0.854 |
| Bone Volume | supinator | trapezius | 5.427 | 0.030 | 0.832 |
| Bone Volume | serratus anterior | supinator | -5.412 | 0.031 | -0.931 |
| Bone Volume | flexor digitorum superficialis | supinator | -5.309 | 0.042 | -0.739 |
| Product | flexor digitorum superficialis | supinator | -5.282 | 0.046 | -0.650 |
| Bone Volume | deltoid | supinator | -5.278 | 0.046 | -0.776 |
| Bone Volume | gemelli | supinator | -5.268 | 0.048 | -0.998 |

*Effect of Sex on Product (Height*Weight) Muscle Volume Normalization*

When analyzing the effect of sex on the error in predicted muscle volume utilizing the sex specific product normalization method's linear regression, there was no statistically significant main or interactions effects. Specifically for the effect of muscle ($F(70, 5530) = .654$, $p_{GG} = 0.849$, $\eta_2 = 0.008$, $\epsilon = 0.249$), sex ($F(1, 79) = 0.061$, $p = 0.805$, $\eta_2 = 0.001$), or interaction ($F(70, 5530) = 0.361$, $p = 1.000$, $\eta_2 = 0.005$).

**Muscle Length Scaling**

When analyzing the effect of sex on the error in predicted muscle length utilizing the bone length normalization linear regression, there was not a statistically significant main effect of muscle ($F(70, 5530) = 1.459$, $p_{GG} = 0.110$, $\eta_2 = 0.018$, $\epsilon = 0.209$), sex ($F(1, 79) = 0.273$, $p = 0.603$, $\eta_2 = 0.003$) or interaction ($F(70, 5530) = 0.292$, $p = 0.999$, $\eta_2 = 0.004$).

### Analysis of Metric Variation Across Muscles and Biologic Sex

Main and interaction effect results of the two-way Mixed ANOVA are reported for each dependent variable below for both individual muscles and functional muscle groups (included for asymmetry, fat infiltration, and normalized volume). Post hoc comparison results with significant findings are reported in **Table S3-8**, where positive t-statistic values indicate the metric was significantly larger in females than males and a negative t-statistic indicates the metric was significantly larger in males than females. For the analysis of ratio of raw volume to expected volume using the bone volume linear regression model, elements of the male and female linear regression models are reported in **Table S9**.

*Asymmetry Magnitude (%)*

For individual muscle structures, there was no statistically significant interaction effect between biologic sex and muscle ($F(70,2450) = 0.99$, $p = 0.503$, $\eta_2 = 0.03$). Additionally, there was no statistically significant main effect of biologic sex ($F(1,35) = 3.67$, $p = 0.063$, $\eta_2 = 0.09$). The main effect of muscle demonstrated a significant difference in asymmetry magnitude between individual muscle structures ($F(70, 2450) = 12.62$, $p_{GG} < 0.001$, $\eta_2 = 0.27$, $\epsilon = 0.25$). Post-hoc comparisons are reported in **Table S3**.

For functional muscle groupings, there was no statistically significant interaction effect between biologic sex and muscle group ($F(18,612) = 1.22$, $p = 0.236$, $\eta_2 = 0.03$). Additionally, there was no statistically significant main effect of biologic sex ($F(1,34) = 1.81$, $p = 0.188$, $\eta_2 = 0.05$). The main effect of muscle group demonstrated a significant difference in asymmetry magnitude between functional muscle groups ($F(18,612) = 22.3$, $p_{GG} < 0.001$, $\eta_2 = 0.4$, $\epsilon = 0.24$). Post-hoc comparisons are reported in **Table S3**.

The muscle with the largest mean asymmetry magnitudes for both males ($\mu = 12.24 \pm 5.95$ %) and females ($\mu = 13.28 \pm 5.93$ %) was the supinator. The muscles with the lowest mean asymmetry magnitudes were the triceps brachii for males ($\mu = 1.55 \pm 1.33\%$) and the rectus abdominis for females ($\mu = 1.55 \pm 1.34\%$).

The most asymmetric functional muscle grouping in both males and females is the wrist pronators ($\mu_{Males} = 7.14 \pm 4.43\%$; $\mu_{Females} = 9.63 \pm 6.28\%$). The least asymmetric functional muscle grouping in males is the hip flexors ($\mu = 1.24 \pm 0.88\%$) and trunk flexors in females ($\mu = 1 \pm 0.65\%$).

*Fat Fraction (%)*

For the fat fraction values, there was no statistically significant main effect of biologic sex for individual muscles ($F(1,42) = 3.33$, $p = 0.075$, $\eta_2 = 0.07$) or functional muscle groups ($F(1,34) = 3.91$, $p = 0.06$, $\eta_2 = 0.1$). The main effect of muscle demonstrated a significant difference in fat fraction between individual muscles ($F(70,2940) = 64.32$, $p_{GG} < 0.001$, $\eta_2 = 0.6$, $\epsilon = 0.09$) as well as functional muscle groups ($F(18,612) = 71.8$, $p_{GG} < 0.001$, $\eta_2 = 0.68$, $\epsilon = 0.218$). There was also a statistically significant interaction effect between biologic sex and individual muscles ($F(70,2940) = 3.16$, $p < 0.001$, $\eta_2 = 0.07$). In functional groups, there was no statistically significant interaction effect ($F(18,612) = 1.21$, p_unc = 0.25, $\eta_2 = 0.03$). Post-hoc comparisons are reported in **Table S4**.

The statistical analysis of the effect of biologic sex and muscle or functional muscle group on fat fraction percentage indicate that normative ranges of fat fraction are muscle/group-specific and sex-specific. Females exhibited higher average intramuscular fat fraction in most muscles, demonstrating a difference in expected tissue composition based on biologic sex. Some muscles showed variations in average fat fraction of multiple percentage points, such as the tensor fasciae latae ($\mu_{Male}$ = 6.77 ± 2.63% and $\mu_{Female}$ = 11.08 ± 10.83%). The muscle with the highest levels of fat fraction, outside of the subclavius ($\mu_{Male}$ = 17.88 ± 6.63% and $\mu_{Female}$ = 20.0 ± 7.19%) was the trapezius in both males and females ($\mu_{Male}$ = 12.93 ± 5.47% and $\mu_{Female}$ =16.84 ± 4.66%). Due to the low absolute raw volume of the subclavius, its lack of adjacency to other muscle tissue, and its orientation in the plane of image acquisition, it is more prone to intermuscular or subcutaneous fat tissue inflating the fat fraction percentage for such a small muscle. The muscle with the lowest level of fat fraction was the flexor digitorum longus across both males and females ($\mu_{Male}$ = 4.37 ± 1.59% and $\mu_{Female}$ = 4.59 ± 0.99%). The functional muscle group with the highest fat fraction is the trunk extensors in males ($\mu$ = 10.03 ± 2.1%) and the scapular stabilizers in females ($\mu$ = 11.99 ± 2.08%). The functional muscle group with the lowest fat fraction in both males and females is the ankle dorsiflexors ($\mu_{Male}$ = 5.01 ± 1.67% and $\mu_{Female}$ = 5.27 ± 1.03%).

*Normalized Muscle Volume ($cm^2$/kg)*

For individual muscle structures, the main effect of biologic sex demonstrated that there were significantly higher normalized volume values in male subjects than female subjects ($F(1,42) = 25.95$, $p < 0.001$, $\eta^2 = 0.38$). The main effect of muscle demonstrated a significant difference in normalized volume values between individual muscle structures ($F(70,2940) = 1342.97$, $p_{GG} < 0.001$, $\eta^2 = 0.97$, $\epsilon = 0.05$). There was also a statistically significant interaction effect between biologic sex and muscle ($F(70, 2940) = 12.3$, $p\_unc < 0.001$, $\eta^2 = 0.23$). Post-hoc comparisons are reported in **Table S5**.

For functional muscle groupings, the main effect of biologic sex demonstrated that there were significantly higher normalized volume values in male subjects than female subjects ($F(1,46) = 21.05$, $p < 0.001$, $\eta^2 = 0.31$). The main effect of muscle demonstrated a significant difference in normalized volume values between functional muscle groups ($F(18,828) = 555.33$, $p_{GG} < 0.001$, $\eta^2 = 0.92$, $\epsilon = 0.29$). There was also a statistically significant interaction effect between biologic sex and muscle ($F(18,828) = 6.11$, $p\_unc < 0.001$, $\eta^2 = 0.12$). Post-hoc comparisons are reported in **Table S5**.

There was an overall trend across nearly all muscles of the mean normalized muscle volume was larger in male subjects than in females and the majority were significantly different. This trend is indicative of more of the male subject size being comprised of muscle volume as compared to females. Muscles that exhibited larger mean normalized volumes in females than males include the soleus, medial gastrocnemius, lateral gastrocnemius, fibulari, phalangeal extensors, piriformis, and rectus abdominis. The soleus, medial gastrocnemius, lateral gastrocnemius, and fibulari contributed to females having a higher mean normalized volume in the ankle plantar flexors, where all other functional groups were higher in males. Overall, the presence of statistically significant normalized muscle volumes at both the individual muscle and functional group level fraction indicate that this metric of muscle development is muscle/group dependent and is sex-specific.

*Individual Muscle Volume Fraction of Total Muscle Volume*

For individual muscles, the main effect of biologic sex demonstrated significant differences between males and females ($F(1,35) = \infty$, $p < 0.001$, $\eta 2 = 1.0$). The main effect of muscle demonstrated significant differences in volume fractions ($F(70,2450) = 2001.06$, $p_{GG} < 0.001$, $\eta 2 = 0.98$, $\epsilon = 0.1$). There was also a statistically significant interaction effect between biologic sex and muscle ($F(70, 2450) = 8.15$, $p < 0.001$, $\eta 2 = 0.19$). Post-hoc comparisons are reported in **Table S6**.

*Actual vs. Expected Muscle Volume Difference – Height and Mass Product Linear Regression*

For individual muscle structures, the main effect of biologic sex demonstrated that there were significantly higher difference between raw and predicted volumes (calculated using the height and mass product linear regression model, elements detailed in **Table 2**) in female subjects than male subjects ($F(1,39) = 9.68$, $p = 0.003$, $\eta 2 = 0.2$). There was no statistically significant main effect of muscle ($F(70,2730) = 0.04$, $p_{GG} = 0.99$, $\eta 2 = 0.001$, $\epsilon = 0.05$). There was a statistically significant interaction effect between biologic sex and muscle ($F(70, 2730) = 4.76$, p_unc < 0.001, $\eta 2 = 0.11$). Post-hoc comparisons are reported in **Table S7**.

There was an overall trend across most muscles where the mean difference between raw volumes and predicted volumes, calculated using the height and mass product linear regression, was lower in female subjects than male. This trend is indicative that the scaling method using the height and mass product linear regression model is sex-specific and requires equation elements generated for each biologic sex separately.

*Actual vs. Expected Muscle Volume Difference – Associated Bone Volume Linear Regression*

For individual muscle structures, there were no statistically significant main effects of either biologic sex ($F(1,39) = 1.68$, $p = 0.203$, $\eta 2 = 0.04$) or muscle ($F(70,2730) = 0.146$, $p_{GG} = 0.963$, $\eta 2 = 0.003$, $\epsilon = 0.06$). Predicted volumes were calculated using the associated bone volume linear regression model, elements reported in **Table S9**. There was also no statistically significant interaction effect between biologic sex and muscle ($F(70, 2730) = 1.2$, p_unc = 0.13, $\eta 2 = 0.03$). Post-hoc comparisons are reported in **Table S8**.

**Table S3:** Post-hoc analysis results of the interaction effect for the dependent variable of asymmetry reported for individual muscles and functional muscle groupings. The contrast was Biologic Sex*Muscle.

| Muscle/Group Names | Mean ± STDEV (%) | | DoF | t-Statistic | p-Value |
|---|---|---|---|---|---|
| | Male | Female | | | |
| adductor brevis | 3.67 ± 3.11 | 3.82 ± 2.53 | 34.8 | 0.44 | 1 |
| adductor longus | 2.45 ± 1.89 | 2.52 ± 2.01 | 34.5 | 0.21 | 1 |
| adductor magnus | 2.41 ± 1.7 | 2.84 ± 2.11 | 27.7 | 1.05 | 1 |
| anconeus | 5.89 ± 5.87 | 6.46 ± 5.46 | 27.3 | 1.35 | 1 |
| biceps brachii | 3.14 ± 2.23 | 4.08 ± 3.42 | 28.6 | 0.33 | 1 |
| biceps femoris: long head | 3.21 ± 1.97 | 3.84 ± 2.6 | 26.4 | 0.9 | 1 |
| biceps femoris: short head | 4.19 ± 2.34 | 5.42 ± 3.01 | 33.5 | 0.32 | 1 |
| brachialis | 2.15 ± 1.13 | 2.78 ± 2.14 | 23.6 | 0.65 | 1 |
| brachioradialis | 4.28 ± 3.71 | 3.43 ± 2.33 | 31.5 | -0.74 | 1 |
| coracobrachialis | 8.03 ± 5.12 | 6.98 ± 7.11 | 29.5 | -1.08 | 1 |
| deep forearm extensors | 3.92 ± 2.94 | 4.58 ± 4.06 | 23.2 | 1.13 | 1 |
| deltoid | 2.08 ± 1.07 | 1.76 ± 1.05 | 34.4 | -1.44 | 1 |
| digit extensors | 3.39 ± 2.13 | 6.38 ± 5.56 | 25 | 1.97 | 1 |
| erector spinae ub | 1.62 ± 1.4 | 2.8 ± 1.83 | 30.6 | 1.91 | 1 |
| extensor carpi radialis | 6.12 ± 3.96 | 5.35 ± 3.37 | 35 | -0.85 | 1 |
| extensor carpi ulnaris | 9.34 ± 4.04 | 10.71 ± 6.3 | 27 | 0.97 | 1 |
| external oblique | 4.27 ± 3.27 | 3.22 ± 2.22 | 32.2 | -1.25 | 1 |
| fibulari | 4.55 ± 3.88 | 6.17 ± 4.33 | 33.8 | 0.85 | 1 |
| flexor carpi radialis and palmaris longus | 8.64 ± 5.95 | 6.99 ± 4.29 | 32.6 | -0.68 | 1 |
| flexor carpi ulnaris | 5.3 ± 3.52 | 6.39 ± 4.5 | 28.4 | 0.57 | 1 |
| flexor digitorum longus | 6.16 ± 5.12 | 6.87 ± 3.83 | 35 | 0.14 | 1 |
| flexor digitorum profundus | 5.55 ± 4.33 | 6.13 ± 4.17 | 34.7 | 0.42 | 1 |
| flexor digitorum superficialis | 3.96 ± 3.64 | 2.89 ± 1.92 | 29.8 | -1.39 | 1 |
| flexor hallucis longus | 4.28 ± 3.8 | 4.83 ± 3.3 | 35 | 0.79 | 1 |
| flexor pollicis longus | 4.72 ± 3.72 | 8.46 ± 8.99 | 18.6 | 1.81 | 1 |
| gastrocnemius: lateral head | 2.7 ± 2.08 | 3.53 ± 3.21 | 28.9 | -0.1 | 1 |
| gastrocnemius: medial head | 3.44 ± 2.42 | 2.82 ± 2.15 | 34.9 | -0.73 | 1 |
| gemelli | 6.09 ± 5.94 | 5.85 ± 4.36 | 34 | -0.62 | 1 |
| gluteus maximus | 2.3 ± 1.94 | 1.88 ± 1.44 | 34.9 | -0.67 | 1 |
| gluteus medius | 2.5 ± 1.99 | 2.82 ± 2.2 | 33.1 | -0.07 | 1 |
| gluteus minimus | 3.99 ± 2.45 | 4.27 ± 2.46 | 34.1 | -0.15 | 1 |
| gracilis | 5.2 ± 3 | 4.06 ± 2.98 | 34.9 | -1.87 | 1 |
| iliacus | 2.67 ± 2.59 | 3.42 ± 2.64 | 32.8 | -0.58 | 1 |
| infraspinatus and teres minor | 3.17 ± 1.83 | 4.44 ± 7.15 | 19.2 | 0.1 | 1 |
| latissimus dorsi | 2.73 ± 2.19 | 4.15 ± 3.18 | 27.5 | 1.2 | 1 |
| levator scapulae | 3.9 ± 2.69 | 7.36 ± 5.25 | 26.1 | 2.38 | 1 |
| multifidus ub | 3.12 ± 2.84 | 3.39 ± 2.89 | 34.3 | -0.43 | 1 |
| obturator externus | 2.34 ± 1.9 | 3.32 ± 3.41 | 26.6 | 1.14 | 1 |
| obturator internus | 5.34 ± 4.31 | 7.2 ± 4.71 | 32.6 | 0.77 | 1 |
| pectineus | 2.46 ± 2.07 | 4.02 ± 3.08 | 26.4 | 1.89 | 1 |
| pectoralis major | 2.46 ± 1.57 | 3.29 ± 2.64 | 29.6 | 0.83 | 1 |
| pectoralis minor | 4.38 ± 4.14 | 5.89 ± 5.84 | 30.6 | 0.93 | 1 |
| phalangeal extensors | 3.39 ± 1.64 | 3.16 ± 3.01 | 26.3 | -0.5 | 1 |
| piriformis | 6.06 ± 4.34 | 6.09 ± 5.47 | 29.5 | 1.11 | 1 |
| popliteus | 3.76 ± 3.36 | 5.5 ± 3.87 | 32 | 0.81 | 1 |
| pronator quadratus | 7.33 ± 4.89 | 9.3 ± 10.44 | 21.7 | 1.09 | 1 |
| pronator teres | 8.65 ± 5.08 | 10.48 ± 8.14 | 26.8 | 0.82 | 1 |
| psoas major | 2.7 ± 1.75 | 3.07 ± 2.1 | 31.6 | 0.66 | 1 |
| quadratus femoris | 5.04 ± 3.59 | 6.24 ± 4.96 | 29.4 | 0.8 | 1 |
| quadratus lumborum | 4.87 ± 4.91 | 4.12 ± 4.23 | 31.9 | -1.55 | 1 |
| rectus abdominis | 2.18 ± 1.93 | 1.57 ± 1.29 | 32.7 | -1.48 | 1 |
| rectus femoris | 2.39 ± 2.19 | 3.94 ± 3.83 | 23.7 | 1.25 | 1 |
| rhomboid | 4.2 ± 2.71 | 4.87 ± 4.06 | 26.4 | 0.42 | 1 |
| sartorius | 2.83 ± 2.04 | 2.76 ± 2.43 | 29.7 | 0.62 | 1 |
| semimembranosus | 2.54 ± 1.85 | 2.22 ± 1.99 | 34.9 | -1.07 | 1 |
| semitendinosus | 2.83 ± 2.08 | 4.07 ± 3.27 | 24.8 | 1.28 | 1 |
| serratus anterior | 3.58 ± 2.8 | 3.59 ± 3.03 | 32.3 | 0.2 | 1 |
| soleus | 2.57 ± 1.86 | 2.73 ± 1.73 | 34.8 | -0.73 | 1 |
| subclavius | 10.09 ± 8.01 | 8.78 ± 5.38 | 33.1 | -0.35 | 1 |
| subscapularis | 5.64 ± 4.29 | 4.44 ± 3.39 | 34.4 | -0.91 | 1 |

| | | | | | |
|---|---|---|---|---|---|
| **supinator** | 12.24 ± 6.08 | 13.28 ± 6.07 | 32.3 | 0.67 | 1 |
| **supraspinatus** | 3.78 ± 3.39 | 3.78 ± 2.43 | 34.4 | -0.32 | 1 |
| **tensor fasciae latae** | 4.44 ± 2.52 | 3.35 ± 3.34 | 26.6 | -0.26 | 1 |
| **teres major** | 6.12 ± 3.57 | 7.61 ± 7.17 | 25.2 | 0.3 | 1 |
| **tibialis anterior** | 2.78 ± 1.93 | 3.19 ± 2.33 | 33.3 | -0.04 | 1 |
| **tibialis posterior** | 4.84 ± 4.68 | 4.36 ± 3.71 | 32.3 | -1.02 | 1 |
| **trapezius** | 3.71 ± 2 | 3.31 ± 2.96 | 26.3 | 0.1 | 1 |
| **triceps brachii** | 1.55 ± 1.37 | 1.7 ± 1.4 | 33.2 | 0.78 | 1 |
| **vastus intermedius** | 3.82 ± 2.57 | 5.04 ± 4.05 | 24.4 | 1.44 | 1 |
| **vastus lateralis** | 2.73 ± 1.55 | 2.39 ± 2.1 | 28.7 | -0.71 | 1 |
| **vastus medialis** | 2.31 ± 1.6 | 3.86 ± 2.94 | 22.7 | 2.49 | 1 |
| **Ankle Dorsiflexors** | 2.53 ± 1.7 | 2.27 ± 1.83 | 32.5 | -0.34 | 1 |
| **Ankle Plantar Flexors** | 1.56 ± 1.22 | 1.67 ± 1.25 | 32.8 | -1.19 | 1 |
| **Elbow Extensors** | 1.61 ± 1.46 | 1.64 ± 1.3 | 34 | 0.49 | 1 |
| **Elbow Flexors** | 2.25 ± 1.85 | 2.57 ± 1.91 | 33.9 | -0.54 | 1 |
| **Hip Abductors** | 2.3 ± 1.77 | 2.53 ± 1.74 | 33.9 | 0.14 | 1 |
| **Hip Adductors** | 1.9 ± 1.32 | 2.1 ± 1.78 | 26.4 | 0.79 | 1 |
| **Hip Extensors** | 1.68 ± 1.21 | 1.32 ± 1.14 | 33.3 | -0.74 | 1 |
| **Hip External Rotators** | 1.84 ± 0.94 | 2.73 ± 2.06 | 20.4 | 1.43 | 1 |
| **Hip Flexors** | 1.24 ± 0.9 | 1.58 ± 1.3 | 26.9 | 1.33 | 1 |
| **Knee Extensors** | 1.55 ± 1 | 2.18 ± 2.3 | 20 | 1.15 | 1 |
| **Knee Flexors** | 1.55 ± 0.97 | 1.76 ± 1.12 | 34 | 0.16 | 1 |
| **Scapular Stabilizers** | 1.39 ± 1.18 | 1.96 ± 1.39 | 27.4 | 1.51 | 1 |
| **Shoulder Abductors** | 1.83 ± 1.2 | 2.14 ± 1.53 | 34 | -0.41 | 1 |
| **Shoulder Adductors** | 2.46 ± 1.84 | 2.71 ± 2.22 | 30.5 | 0.12 | 1 |
| **Trunk Extensors** | 1.31 ± 1.05 | 1.93 ± 1.56 | 30.2 | 1.35 | 1 |
| **Trunk Flexors** | 2.18 ± 1.6 | 1 ± 0.67 | 24.4 | -2.84 | 0.17 |
| **Wrist Extensors** | 2.35 ± 2.29 | 3.17 ± 2.1 | 33.7 | 0.85 | 1 |
| **Wrist Flexors** | 3.85 ± 2.43 | 4.18 ± 2.34 | 34 | 0.32 | 1 |
| **Wrist Pronators** | 7.14 ± 4.53 | 9.63 ± 6.43 | 30.2 | 1.33 | 1 |

**Table S4:** Post-hoc analysis results of the interaction effect for the dependent variable of fat infiltration reported for individual muscles and functional muscle groupings. The contrast was Biologic Sex*Muscle.

| Muscle/Group Names | Mean ± STDEV (%) | | DoF | t-Statistic | p-Value |
|---|---|---|---|---|---|
| | Male | Female | | | |
| adductor brevis | 4.95 ± 1.68 | 5.18 ± 1 | 33.4 | 0.43 | 1 |
| adductor longus | 4.7 ± 2.24 | 5.03 ± 1.35 | 27.4 | 0.36 | 1 |
| adductor magnus | 6.1 ± 1.92 | 6.48 ± 2.02 | 41.7 | 0.49 | 1 |
| anconeus | 5.42 ± 1.81 | 7.42 ± 1.87 | 42 | 4.35 | 0.006 |
| biceps brachii | 6.21 ± 1.2 | 6.84 ± 1.73 | 32.8 | 1.48 | 1 |
| biceps femoris: long head | 5.7 ± 2.2 | 6.44 ± 2.22 | 40.8 | 1.28 | 1 |
| biceps femoris: short head | 6.22 ± 1.81 | 7.59 ± 2.15 | 39.5 | 2.34 | 1 |
| brachialis | 5.96 ± 1.03 | 6.34 ± 1.38 | 34.5 | 0.65 | 1 |
| brachioradialis | 6.02 ± 1.71 | 6.91 ± 1.57 | 41.4 | 2.33 | 1 |
| coracobrachialis | 8.23 ± 2.12 | 8.39 ± 1.72 | 39.1 | -0.14 | 1 |
| deep forearm extensors | 6.99 ± 1.49 | 7.13 ± 1.81 | 36.4 | 0.02 | 1 |
| deltoid | 9.84 ± 2.84 | 10.88 ± 2.05 | 38.3 | 1.74 | 1 |
| digit extensors | 8.03 ± 2.93 | 8.85 ± 2.14 | 38.1 | 1.13 | 1 |
| erector spinae ub | 9.98 ± 2.28 | 11.33 ± 2.71 | 36.7 | 1.84 | 1 |
| extensor carpi radialis | 6.07 ± 1.57 | 6.74 ± 1.58 | 41.8 | 1.78 | 1 |
| extensor carpi ulnaris | 8.22 ± 5.49 | 9.43 ± 3.61 | 37.3 | 1.22 | 1 |
| external oblique | 8.36 ± 3.51 | 10.43 ± 6.14 | 31.4 | 1.92 | 1 |
| fibulari | 6.92 ± 2.79 | 7.26 ± 1.9 | 34.4 | 0.72 | 1 |
| flexor carpi radialis and palmaris longus | 6.19 ± 1.5 | 6.75 ± 1.86 | 42 | 1.63 | 1 |
| flexor carpi ulnaris | 7.37 ± 2.23 | 9.08 ± 2.99 | 41.9 | 1.99 | 1 |
| flexor digitorum longus | 4.37 ± 1.61 | 4.59 ± 1 | 32.3 | 0.82 | 1 |
| flexor digitorum profundus | 6.18 ± 1.14 | 6.62 ± 1.64 | 41.1 | 0.61 | 1 |
| flexor digitorum superficialis | 6.56 ± 1.53 | 7.11 ± 1.39 | 41.1 | 1.38 | 1 |
| flexor hallucis longus | 5.61 ± 3.45 | 5.24 ± 1.1 | 26.6 | -0.6 | 1 |
| flexor pollicis longus | 5.73 ± 1.76 | 5.52 ± 1.47 | 40.3 | -0.94 | 1 |
| gastrocnemius: lateral head | 5.95 ± 1.44 | 6.05 ± 1.49 | 40.9 | 0.41 | 1 |
| gastrocnemius: medial head | 5.37 ± 1.23 | 5.76 ± 2 | 31.8 | 1 | 1 |
| gemelli | 9.26 ± 2.76 | 10.72 ± 3.15 | 39.7 | 1.64 | 1 |
| gluteus maximus | 9.41 ± 3.49 | 11.69 ± 5.89 | 31.8 | 1.74 | 1 |
| gluteus medius | 7.97 ± 2.3 | 7.81 ± 2.48 | 40.9 | 0.16 | 1 |
| gluteus minimus | 8.2 ± 2.51 | 8.28 ± 1.8 | 36.3 | 0.21 | 1 |
| gracilis | 4.67 ± 2.19 | 7.23 ± 2.52 | 40 | 3.95 | 0.021 |
| iliacus | 7.48 ± 1.49 | 7.31 ± 1.63 | 41.2 | 0.19 | 1 |
| infraspinatus and teres minor | 6.84 ± 1.55 | 7.25 ± 1.29 | 41.6 | 0.89 | 1 |
| latissimus dorsi | 9.28 ± 2.2 | 11.16 ± 3.69 | 30.5 | 2.12 | 1 |
| levator scapulae | 7.49 ± 3.84 | 8.52 ± 2.55 | 40.4 | 1.13 | 1 |
| multifidus ub | 10.6 ± 2.28 | 11.87 ± 2.96 | 36.4 | 1.58 | 1 |
| obturator externus | 8.43 ± 2.58 | 8.2 ± 1.72 | 38.5 | -0.13 | 1 |
| obturator internus | 7.15 ± 2.21 | 6.71 ± 1.81 | 37.5 | -0.2 | 1 |
| pectineus | 5.88 ± 1.77 | 6.15 ± 1.3 | 39.7 | 0.85 | 1 |
| pectoralis major | 9.41 ± 2.48 | 10.16 ± 2.34 | 42 | 0.76 | 1 |
| pectoralis minor | 8.61 ± 2.64 | 9.38 ± 2.28 | 41.6 | 1.01 | 1 |
| phalangeal extensors | 5.14 ± 2.2 | 5.3 ± 1.33 | 36.4 | 0.57 | 1 |
| piriformis | 11 ± 3.27 | 9.79 ± 2.15 | 37 | -1.31 | 1 |
| popliteus | 5.88 ± 1.49 | 5.63 ± 1.47 | 41.3 | -0.41 | 1 |
| pronator quadratus | 11.27 ± 2.8 | 11.03 ± 4.12 | 38.3 | -0.31 | 1 |
| pronator teres | 7.81 ± 2.34 | 8.43 ± 1.61 | 40.5 | 1.21 | 1 |
| psoas major | 8.88 ± 2.34 | 8.33 ± 1.61 | 38.8 | -0.79 | 1 |
| quadratus femoris | 6.72 ± 1.92 | 7.83 ± 3.24 | 31.3 | 1.57 | 1 |
| quadratus lumborum | 8.55 ± 2.46 | 10.83 ± 3.27 | 37.2 | 2.99 | 0.336 |
| rectus abdominis | 11.99 ± 3.96 | 11.31 ± 4.39 | 40.3 | -0.33 | 1 |
| rectus femoris | 4.9 ± 1.64 | 6.03 ± 1.46 | 40.8 | 2.81 | 0.495 |
| rhomboid | 7.39 ± 2.02 | 8.73 ± 2.29 | 41.1 | 2.27 | 1 |
| sartorius | 5.69 ± 1.88 | 7.83 ± 3.1 | 30.3 | 3.03 | 0.336 |
| semimembranosus | 6.38 ± 2.21 | 6.64 ± 2.44 | 40.4 | 0.29 | 1 |
| semitendinosus | 5.38 ± 1.51 | 6.15 ± 2.1 | 36.6 | 1.37 | 1 |
| serratus anterior | 7.94 ± 2.37 | 9.89 ± 3.58 | 31.8 | 2.24 | 1 |
| soleus | 5.59 ± 2.11 | 5.24 ± 1.5 | 38.8 | -0.64 | 1 |
| subclavius | 17.88 ± 6.7 | 19.77 ± 7.36 | 40.8 | 1.19 | 1 |
| subscapularis | 6.16 ± 1.35 | 6.56 ± 1.06 | 40.2 | 0.77 | 1 |

| | | | | | |
|---|---|---|---|---|---|
| **supinator** | 8.6 ± 1.83 | 8.32 ± 2.17 | 38.8 | -0.54 | 1 |
| **supraspinatus** | 8.9 ± 3 | 11.36 ± 3.17 | 41.3 | 2.58 | 0.877 |
| **tensor fasciae latae** | 6.77 ± 2.66 | 11.08 ± 10.95 | 22.4 | 2.19 | 1 |
| **teres major** | 8.01 ± 2.33 | 8.23 ± 1.75 | 35.1 | 0.25 | 1 |
| **tibialis anterior** | 4.91 ± 1.52 | 5.23 ± 0.9 | 34.2 | 1.11 | 1 |
| **tibialis posterior** | 4.97 ± 1.55 | 4.93 ± 0.72 | 28.4 | -0.24 | 1 |
| **trapezius** | 12.93 ± 5.53 | 16.94 ± 4.76 | 41.3 | 2.86 | 0.449 |
| **triceps brachii** | 6.6 ± 1.02 | 7.11 ± 1.59 | 31.2 | 1.05 | 1 |
| **vastus intermedius** | 4.69 ± 1.39 | 5.16 ± 1.25 | 38.5 | 1.21 | 1 |
| **vastus lateralis** | 5.56 ± 1.59 | 6.27 ± 1.47 | 41.9 | 1.8 | 1 |
| **vastus medialis** | 4.55 ± 1.22 | 5.12 ± 1.12 | 41.8 | 1.91 | 1 |
| **Ankle Dorsiflexors** | 5.01 ± 1.69 | 5.27 ± 1.04 | 22.5 | 1.66 | 1 |
| **Ankle Plantar Flexors** | 5.63 ± 1.65 | 5.65 ± 1.43 | 26.1 | 0.3 | 1 |
| **Elbow Extensors** | 6.65 ± 0.99 | 7.09 ± 1.5 | 34 | 1.32 | 1 |
| **Elbow Flexors** | 6.08 ± 1.08 | 6.68 ± 1.46 | 33 | 2.19 | 0.576 |
| **Hip Abductors** | 7.86 ± 2.28 | 8.31 ± 3.04 | 32.9 | 1.01 | 1 |
| **Hip Adductors** | 5.57 ± 1.67 | 6.16 ± 1.72 | 31.8 | 1.49 | 1 |
| **Hip Extensors** | 7.92 ± 2.78 | 9.59 ± 4.34 | 34 | 2.06 | 0.658 |
| **Hip External Rotators** | 8.67 ± 1.96 | 8.74 ± 1.74 | 32.3 | 0.59 | 1 |
| **Hip Flexors** | 7.48 ± 1.48 | 7.77 ± 1.82 | 34 | 1.25 | 1 |
| **Knee Extensors** | 5.1 ± 1.38 | 5.8 ± 1.26 | 29.1 | 2.84 | 0.156 |
| **Knee Flexors** | 5.76 ± 1.58 | 6.34 ± 1.94 | 32.2 | 1.34 | 1 |
| **Scapular Stabilizers** | 9.98 ± 3.24 | 11.99 ± 2.11 | 29.2 | 2.12 | 0.634 |
| **Shoulder Abductors** | 8.98 ± 2.4 | 10.01 ± 1.79 | 24.9 | 1.51 | 1 |
| **Shoulder Adductors** | 8.77 ± 1.92 | 10.09 ± 2.41 | 31 | 1.5 | 1 |
| **Trunk Extensors** | 10.03 ± 2.12 | 11.07 ± 2.02 | 34 | 1.47 | 1 |
| **Trunk Flexors** | 9.77 ± 2.88 | 10.34 ± 4.14 | 33.3 | 0.87 | 1 |
| **Wrist Extensors** | 6.83 ± 1.32 | 7.69 ± 1.45 | 33.2 | 2.54 | 0.287 |
| **Wrist Flexors** | 6.33 ± 1.11 | 7.05 ± 1.49 | 30 | 2.43 | 0.365 |
| **Wrist Pronators** | 8.45 ± 2.08 | 8.85 ± 1.69 | 32.3 | 0.38 | 1 |

**Table S5:** Post-hoc analysis results of the interaction effect for the dependent variable of muscle volume normalized by the height*mass reported for individual muscles and functional muscle groupings. The contrast was Biologic Sex*Muscle.

| Muscle/Group Names | Mean ± STDEV | | DoF | t-Statistic | p-Value |
|---|---|---|---|---|---|
| | Male | Female | | | |
| adductor brevis | 0.008 ± 0.0012 | 0.0078 ± 0.0011 | 41.5 | -0.97 | 1 |
| adductor longus | 0.0141 ± 0.0021 | 0.0127 ± 0.0021 | 42 | -2.6 | 0.304 |
| adductor magnus | 0.0506 ± 0.0099 | 0.0459 ± 0.0082 | 42 | -1.72 | 1 |
| anconeus | 0.0008 ± 0.0002 | 0.0007 ± 0.0001 | 42 | -5.27 | < 0.001 |
| biceps brachii | 0.0149 ± 0.0028 | 0.0105 ± 0.0017 | 36.9 | -7 | < 0.001 |
| biceps femoris: long head | 0.017 ± 0.0027 | 0.0159 ± 0.003 | 41.9 | -2.25 | 0.683 |
| biceps femoris: short head | 0.0081 ± 0.0012 | 0.0068 ± 0.0013 | 41.7 | -4.68 | 0.001 |
| brachialis | 0.0114 ± 0.0017 | 0.0095 ± 0.0015 | 40.9 | -4.88 | 0.001 |
| brachioradialis | 0.006 ± 0.0011 | 0.0041 ± 0.0008 | 40.8 | -7.09 | < 0.001 |
| coracobrachialis | 0.0026 ± 0.0004 | 0.0021 ± 0.0005 | 37.5 | -4.21 | 0.007 |
| deep forearm extensors | 0.0027 ± 0.0006 | 0.0023 ± 0.0004 | 41.8 | -3.28 | 0.076 |
| deltoid | 0.0367 ± 0.0052 | 0.0287 ± 0.0043 | 40.7 | -6.51 | < 0.001 |
| digit extensors | 0.0033 ± 0.0009 | 0.0029 ± 0.0004 | 29.8 | -2.13 | 0.881 |
| erector spinae ub | 0.041 ± 0.0055 | 0.0347 ± 0.0055 | 39.7 | -3.41 | 0.057 |
| extensor carpi radialis | 0.0051 ± 0.0008 | 0.004 ± 0.0006 | 41.8 | -6.38 | < 0.001 |
| extensor carpi ulnaris | 0.0016 ± 0.0005 | 0.0013 ± 0.0004 | 42 | -2.17 | 0.794 |
| external oblique | 0.015 ± 0.002 | 0.0142 ± 0.0031 | 29.2 | -1.12 | 1 |
| fibulari | 0.0095 ± 0.0015 | 0.01 ± 0.0016 | 40.4 | 0.71 | 1 |
| flexor carpi radialis and palmaris longus | 0.0044 ± 0.0011 | 0.0038 ± 0.0006 | 39.7 | -3.1 | 0.116 |
| flexor carpi ulnaris | 0.0033 ± 0.0008 | 0.0027 ± 0.0008 | 38.6 | -3.34 | 0.069 |
| flexor digitorum longus | 0.0019 ± 0.0004 | 0.0017 ± 0.0003 | 42 | -2.93 | 0.155 |
| flexor digitorum profundus | 0.0072 ± 0.0015 | 0.0058 ± 0.0016 | 35.3 | -3.66 | 0.033 |
| flexor digitorum superficialis | 0.0056 ± 0.0012 | 0.0044 ± 0.0007 | 39 | -5.09 | 0.001 |
| flexor hallucis longus | 0.0057 ± 0.0008 | 0.0051 ± 0.0008 | 40.5 | -2.67 | 0.27 |
| flexor pollicis longus | 0.0014 ± 0.0003 | 0.0012 ± 0.0002 | 39.9 | -4.2 | 0.006 |
| gastrocnemius: lateral head | 0.0114 ± 0.0018 | 0.0119 ± 0.0023 | 36.1 | 0.88 | 1 |
| gastrocnemius: medial head | 0.0196 ± 0.0039 | 0.0212 ± 0.0034 | 42 | 1.35 | 1 |
| gemelli | 0.0015 ± 0.0003 | 0.0014 ± 0.0003 | 39.7 | -1.4 | 1 |
| gluteus maximus | 0.0781 ± 0.0108 | 0.0733 ± 0.0087 | 42 | -1.44 | 1 |
| gluteus medius | 0.0284 ± 0.0036 | 0.0271 ± 0.0033 | 41.7 | -2.04 | 0.955 |
| gluteus minimus | 0.0081 ± 0.0011 | 0.0077 ± 0.001 | 40.8 | -2 | 0.984 |
| gracilis | 0.0082 ± 0.0023 | 0.0074 ± 0.0012 | 35.5 | -1.51 | 1 |
| iliacus | 0.0149 ± 0.0016 | 0.0125 ± 0.0018 | 40.9 | -5.77 | < 0.001 |
| infraspinatus and teres minor | 0.0147 ± 0.0018 | 0.0118 ± 0.002 | 41.9 | -6.69 | < 0.001 |
| latissimus dorsi | 0.0367 ± 0.0059 | 0.0285 ± 0.0058 | 40.7 | -4.57 | 0.002 |
| levator scapulae | 0.004 ± 0.0006 | 0.003 ± 0.001 | 41.6 | -5.59 | < 0.001 |
| multifidus ub | 0.0167 ± 0.0019 | 0.0145 ± 0.0036 | 41.6 | -3 | 0.136 |
| obturator externus | 0.0038 ± 0.0008 | 0.0036 ± 0.0006 | 41 | -1.7 | 1 |
| obturator internus | 0.0018 ± 0.0004 | 0.0016 ± 0.0004 | 42 | -2.8 | 0.206 |
| pectineus | 0.0052 ± 0.0007 | 0.0042 ± 0.0005 | 39.8 | -6.17 | < 0.001 |
| pectoralis major | 0.0315 ± 0.0055 | 0.0213 ± 0.0048 | 42 | -7.02 | < 0.001 |
| pectoralis minor | 0.0041 ± 0.0006 | 0.0032 ± 0.0007 | 37.1 | -5.71 | < 0.001 |
| phalangeal extensors | 0.0068 ± 0.0007 | 0.007 ± 0.0012 | 29.5 | 0.56 | 1 |
| piriformis | 0.0033 ± 0.0007 | 0.0036 ± 0.0009 | 36.7 | 1.08 | 1 |
| popliteus | 0.0016 ± 0.0003 | 0.0014 ± 0.0002 | 38.9 | -3.04 | 0.131 |
| pronator quadratus | 0.0007 ± 0.0002 | 0.0006 ± 0.0001 | 42 | -2.69 | 0.262 |
| pronator teres | 0.0031 ± 0.0005 | 0.0026 ± 0.0005 | 39.6 | -4.25 | 0.006 |
| psoas major | 0.0229 ± 0.0033 | 0.0177 ± 0.0027 | 41.9 | -6.38 | < 0.001 |
| quadratus femoris | 0.0023 ± 0.0005 | 0.0019 ± 0.0007 | 40.9 | -3.07 | 0.12 |
| quadratus lumborum | 0.005 ± 0.0008 | 0.0043 ± 0.0007 | 41.3 | -3.66 | 0.029 |
| rectus abdominis | 0.0199 ± 0.0031 | 0.0202 ± 0.0038 | 35.7 | 0.57 | 1 |
| rectus femoris | 0.0216 ± 0.0031 | 0.0189 ± 0.0031 | 41.2 | -3.49 | 0.045 |
| rhomboid | 0.0065 ± 0.0008 | 0.0047 ± 0.0008 | 41.3 | -8.54 | < 0.001 |
| sartorius | 0.0129 ± 0.0016 | 0.0112 ± 0.0019 | 34.7 | -3.17 | 0.107 |
| semimembranosus | 0.0198 ± 0.003 | 0.0187 ± 0.0032 | 40.8 | -1.23 | 1 |
| semitendinosus | 0.0156 ± 0.0041 | 0.0142 ± 0.0022 | 37.1 | -1.38 | 1 |
| serratus anterior | 0.015 ± 0.0017 | 0.0122 ± 0.0016 | 41.6 | -6.64 | < 0.001 |
| soleus | 0.0339 ± 0.0047 | 0.036 ± 0.004 | 42 | 1.49 | 1 |
| subclavius | 0.0006 ± 0.0002 | 0.0005 ± 0.0001 | 35.5 | -4.62 | 0.002 |

| | | | | | |
|---|---|---|---|---|---|
| **subscapularis** | 0.0117 ± 0.0015 | 0.0095 ± 0.0018 | 41.9 | -7.43 | < 0.001 |
| **supinator** | 0.0014 ± 0.0003 | 0.0011 ± 0.0003 | 39.7 | -5.34 | < 0.001 |
| **supraspinatus** | 0.0057 ± 0.0008 | 0.0046 ± 0.0009 | 39.3 | -5.06 | 0.001 |
| **tensor fasciae latae** | 0.0063 ± 0.0013 | 0.0052 ± 0.0011 | 42 | -3.23 | 0.085 |
| **teres major** | 0.0024 ± 0.0005 | 0.0019 ± 0.0004 | 41.1 | -4.55 | 0.002 |
| **tibialis anterior** | 0.009 ± 0.0009 | 0.0091 ± 0.0012 | 37.8 | -0.38 | 1 |
| **tibialis posterior** | 0.0072 ± 0.0013 | 0.0072 ± 0.0014 | 38.9 | -0.44 | 1 |
| **trapezius** | 0.0199 ± 0.0025 | 0.015 ± 0.0041 | 40.4 | -5.45 | < 0.001 |
| **triceps brachii** | 0.0362 ± 0.0063 | 0.0279 ± 0.0055 | 41.2 | -5.49 | < 0.001 |
| **vastus intermedius** | 0.0221 ± 0.0042 | 0.0189 ± 0.0027 | 38.3 | -2.99 | 0.14 |
| **vastus lateralis** | 0.0796 ± 0.0119 | 0.0669 ± 0.0091 | 40.3 | -5.02 | 0.001 |
| **vastus medialis** | 0.041 ± 0.006 | 0.0341 ± 0.0044 | 39.7 | -4.95 | 0.001 |
| **Ankle Dorsiflexors** | 0.0158 ± 0.0015 | 0.0161 ± 0.0021 | 46 | 0.58 | 1 |
| **Ankle Plantar Flexors** | 0.0892 ± 0.0101 | 0.093 ± 0.0101 | 46 | 1.34 | 0.992 |
| **Elbow Extensors** | 0.0385 ± 0.0065 | 0.0274 ± 0.01 | 46 | -4.52 | 0.001 |
| **Elbow Flexors** | 0.0324 ± 0.0049 | 0.0231 ± 0.006 | 46 | -5.86 | < 0.001 |
| **Hip Abductors** | 0.0428 ± 0.0049 | 0.0399 ± 0.004 | 46 | -2.3 | 0.23 |
| **Hip Adductors** | 0.0776 ± 0.0206 | 0.0738 ± 0.0105 | 46 | -0.79 | 1 |
| **Hip Extensors** | 0.1253 ± 0.0307 | 0.1221 ± 0.0131 | 46 | -0.47 | 1 |
| **Hip External Rotators** | 0.0127 ± 0.0019 | 0.0117 ± 0.0031 | 46 | -1.41 | 0.992 |
| **Hip Flexors** | 0.0559 ± 0.0052 | 0.0438 ± 0.0105 | 46 | -5.05 | < 0.001 |
| **Knee Extensors** | 0.1643 ± 0.0221 | 0.1387 ± 0.0159 | 46 | -4.65 | < 0.001 |
| **Knee Flexors** | 0.0896 ± 0.0218 | 0.086 ± 0.0207 | 46 | -0.58 | 1 |
| **Scapular Stabilizers** | 0.0501 ± 0.0049 | 0.0329 ± 0.0155 | 46 | -5.14 | < 0.001 |
| **Shoulder Abductors** | 0.0571 ± 0.0073 | 0.0432 ± 0.0112 | 46 | -5.1 | < 0.001 |
| **Shoulder Adductors** | 0.0813 ± 0.0212 | 0.0575 ± 0.0205 | 46 | -3.94 | 0.004 |
| **Trunk Extensors** | 0.0626 ± 0.0075 | 0.0451 ± 0.0215 | 46 | -3.76 | 0.006 |
| **Trunk Flexors** | 0.0552 ± 0.0134 | 0.0479 ± 0.0163 | 46 | -1.68 | 0.704 |
| **Wrist Extensors** | 0.0119 ± 0.004 | 0.0093 ± 0.0037 | 46 | -2.31 | 0.23 |
| **Wrist Flexors** | 0.0215 ± 0.0055 | 0.016 ± 0.0065 | 46 | -3.18 | 0.029 |
| **Wrist Pronators** | 0.0036 ± 0.0009 | 0.0029 ± 0.001 | 46 | -2.67 | 0.106 |

**Table S6:** Post-hoc analysis results of the interaction effect for the dependent variable of muscle volume fraction of total muscle volume reported for individual muscles and functional muscle groupings. The contrast was Biologic Sex*Muscle.

| Muscle/Group Names | Mean ± STDEV | | DoF | t-Statistic | p-Value |
|---|---|---|---|---|---|
| | Male | Female | | | |
| adductor brevis | 0.0079 ± 0.0009 | 0.0088 ± 0.001 | 32.9 | 3.13 | 0.198 |
| adductor longus | 0.0142 ± 0.0016 | 0.0143 ± 0.0016 | 34 | 0.08 | 1 |
| adductor magnus | 0.0486 ± 0.0045 | 0.0526 ± 0.0064 | 28.4 | 2.16 | 1 |
| anconeus | 0.0008 ± 0.0001 | 0.0008 ± 0.0001 | 34.7 | -2.12 | 1 |
| biceps brachii | 0.0147 ± 0.0017 | 0.0122 ± 0.0011 | 33.3 | -5.48 | < 0.001 |
| biceps femoris: long head | 0.0169 ± 0.0019 | 0.0174 ± 0.0022 | 32.2 | 0.72 | 1 |
| biceps femoris: short head | 0.008 ± 0.001 | 0.0075 ± 0.001 | 34.4 | -1.43 | 1 |
| brachialis | 0.0113 ± 0.0013 | 0.0108 ± 0.001 | 34.8 | -1.39 | 1 |
| brachioradialis | 0.0061 ± 0.0008 | 0.0049 ± 0.0007 | 34.5 | -4.75 | 0.002 |
| coracobrachialis | 0.0026 ± 0.0003 | 0.0024 ± 0.0004 | 30.2 | -1.71 | 1 |
| deep forearm extensors | 0.0028 ± 0.0004 | 0.0026 ± 0.0004 | 34 | -1.38 | 1 |
| deltoid | 0.0362 ± 0.0019 | 0.0326 ± 0.002 | 33.1 | -5.4 | < 0.001 |
| digit extensors | 0.0034 ± 0.0006 | 0.0034 ± 0.0002 | 25.2 | -0.23 | 1 |
| erector spinae ub | 0.0406 ± 0.0043 | 0.0408 ± 0.0047 | 32.8 | 0.1 | 1 |
| extensor carpi radialis | 0.0051 ± 0.0005 | 0.0046 ± 0.0004 | 35 | -3.33 | 0.116 |
| extensor carpi ulnaris | 0.0016 ± 0.0003 | 0.0016 ± 0.0002 | 34.4 | -0.2 | 1 |
| external oblique | 0.015 ± 0.0015 | 0.0158 ± 0.0017 | 32.5 | 1.64 | 1 |
| fibulari | 0.0098 ± 0.0012 | 0.0112 ± 0.0011 | 34.6 | 3.72 | 0.041 |
| flexor carpi radialis and palmaris longus | 0.0045 ± 0.0006 | 0.0043 ± 0.0005 | 35 | -1.09 | 1 |
| flexor carpi ulnaris | 0.0034 ± 0.0004 | 0.0032 ± 0.0004 | 33.8 | -1.83 | 1 |
| flexor digitorum longus | 0.0019 ± 0.0003 | 0.0019 ± 0.0003 | 33.7 | 0.42 | 1 |
| flexor digitorum profundus | 0.0075 ± 0.0008 | 0.0069 ± 0.0007 | 35 | -2.17 | 1 |
| flexor digitorum superficialis | 0.0058 ± 0.0007 | 0.005 ± 0.0005 | 33.7 | -3.72 | 0.042 |
| flexor hallucis longus | 0.0058 ± 0.0007 | 0.006 ± 0.0008 | 31.9 | 1.01 | 1 |
| flexor pollicis longus | 0.0014 ± 0.0002 | 0.0013 ± 0.0002 | 34.2 | -1.28 | 1 |
| gastrocnemius: lateral head | 0.0118 ± 0.0015 | 0.0136 ± 0.0023 | 26.3 | 2.83 | 0.424 |
| gastrocnemius: medial head | 0.0199 ± 0.0033 | 0.0245 ± 0.0032 | 34.6 | 4.28 | 0.009 |
| gemelli | 0.0015 ± 0.0003 | 0.0016 ± 0.0002 | 34.9 | 1.28 | 1 |
| gluteus maximus | 0.0773 ± 0.007 | 0.0854 ± 0.0058 | 35 | 3.88 | 0.027 |
| gluteus medius | 0.0287 ± 0.0027 | 0.0303 ± 0.0032 | 31.7 | 1.57 | 1 |
| gluteus minimus | 0.0081 ± 0.0009 | 0.0089 ± 0.001 | 33.1 | 2.32 | 1 |
| gracilis | 0.0085 ± 0.0011 | 0.0084 ± 0.0011 | 34.4 | -0.5 | 1 |
| iliacus | 0.015 ± 0.0017 | 0.0143 ± 0.0017 | 34 | -1.28 | 1 |
| infraspinatus and teres minor | 0.0147 ± 0.0009 | 0.0136 ± 0.0008 | 34.6 | -4.01 | 0.019 |
| latissimus dorsi | 0.0361 ± 0.0032 | 0.0335 ± 0.0041 | 29.9 | -2.16 | 1 |
| levator scapulae | 0.0041 ± 0.0005 | 0.0036 ± 0.0005 | 34.6 | -2.78 | 0.424 |
| multifidus ub | 0.0168 ± 0.0016 | 0.0176 ± 0.0018 | 32.1 | 1.57 | 1 |
| obturator externus | 0.0038 ± 0.0006 | 0.004 ± 0.0006 | 33.4 | 1.29 | 1 |
| obturator internus | 0.0018 ± 0.0003 | 0.0017 ± 0.0003 | 31.8 | -0.26 | 1 |
| pectineus | 0.0052 ± 0.0007 | 0.0047 ± 0.0004 | 30.9 | -2.88 | 0.365 |
| pectoralis major | 0.0309 ± 0.003 | 0.0244 ± 0.0037 | 30.6 | -5.76 | < 0.001 |
| pectoralis minor | 0.0041 ± 0.0005 | 0.0036 ± 0.0006 | 29.2 | -2.96 | 0.317 |
| phalangeal extensors | 0.007 ± 0.0008 | 0.0082 ± 0.0011 | 28.9 | 3.99 | 0.025 |
| piriformis | 0.0034 ± 0.0004 | 0.0041 ± 0.0011 | 20.4 | 2.39 | 1 |
| popliteus | 0.0016 ± 0.0003 | 0.0016 ± 0.0002 | 32.5 | 0.27 | 1 |
| pronator quadratus | 0.0007 ± 0.0001 | 0.0007 ± 0.0002 | 31.6 | -0.6 | 1 |
| pronator teres | 0.0031 ± 0.0004 | 0.003 ± 0.0004 | 33.3 | -1.03 | 1 |
| psoas major | 0.023 ± 0.0026 | 0.0205 ± 0.0032 | 31 | -2.57 | 0.683 |
| quadratus femoris | 0.0023 ± 0.0004 | 0.0022 ± 0.0005 | 28.8 | -0.75 | 1 |
| quadratus lumborum | 0.0051 ± 0.0006 | 0.0049 ± 0.0007 | 33.1 | -0.98 | 1 |
| rectus abdominis | 0.0196 ± 0.0024 | 0.0231 ± 0.0023 | 34.4 | 4.45 | 0.006 |
| rectus femoris | 0.0216 ± 0.0032 | 0.0214 ± 0.0023 | 34.2 | -0.22 | 1 |
| rhomboid | 0.0066 ± 0.0009 | 0.0054 ± 0.0007 | 34.9 | -4.63 | 0.003 |
| sartorius | 0.013 ± 0.002 | 0.0128 ± 0.002 | 34.4 | -0.26 | 1 |
| semimembranosus | 0.0194 ± 0.0019 | 0.0217 ± 0.003 | 26 | 2.7 | 0.562 |
| semitendinosus | 0.0162 ± 0.0019 | 0.0161 ± 0.0017 | 35 | -0.18 | 1 |
| serratus anterior | 0.015 ± 0.0011 | 0.0138 ± 0.0014 | 30.7 | -3.02 | 0.271 |
| soleus | 0.0352 ± 0.0042 | 0.0413 ± 0.0042 | 34.1 | 4.38 | 0.007 |
| subclavius | 0.0007 ± 0.0002 | 0.0005 ± 0.0001 | 28.7 | -3.34 | 0.128 |

| | | | | | |
|---|---|---|---|---|---|
| **subscapularis** | 0.0118 ± 0.001 | 0.0106 ± 0.0014 | 27.7 | -2.84 | 0.413 |
| **supinator** | 0.0014 ± 0.0002 | 0.0013 ± 0.0002 | 34.5 | -2.52 | 0.721 |
| **supraspinatus** | 0.0057 ± 0.0004 | 0.0054 ± 0.0007 | 25.6 | -1.47 | 1 |
| **tensor fasciae latae** | 0.006 ± 0.0008 | 0.006 ± 0.0012 | 27.1 | -0.06 | 1 |
| **teres major** | 0.0024 ± 0.0004 | 0.0022 ± 0.0004 | 34.3 | -1.95 | 1 |
| **tibialis anterior** | 0.009 ± 0.0011 | 0.0103 ± 0.0008 | 33.7 | 4.12 | 0.015 |
| **tibialis posterior** | 0.0073 ± 0.001 | 0.0085 ± 0.0016 | 27 | 2.69 | 0.562 |
| **trapezius** | 0.0198 ± 0.0013 | 0.0185 ± 0.0019 | 28.4 | -2.33 | 1 |
| **triceps brachii** | 0.0358 ± 0.0032 | 0.0318 ± 0.0036 | 32.4 | -3.52 | 0.074 |
| **vastus intermedius** | 0.0218 ± 0.0029 | 0.0222 ± 0.0017 | 31.3 | 0.46 | 1 |
| **vastus lateralis** | 0.08 ± 0.0079 | 0.0761 ± 0.0082 | 33.6 | -1.46 | 1 |
| **vastus medialis** | 0.041 ± 0.0037 | 0.039 ± 0.0034 | 34.8 | -1.74 | 1 |

**Table S7:** Post-hoc analysis results of the interaction effect for the dependent variable of actual versus expected volume difference using the product of height and mass linear regression model reported for individual muscles. The contrast was Biologic Sex*Muscle.

| Muscle/Group Names | Mean ± STDEV (mL) | | DoF | t-Statistic | p-Value |
|---|---|---|---|---|---|
| | Male | Female | | | |
| adductor brevis | 0.86 ± 17.19 | -0.86 ± 11.92 | 34.1 | -0.68 | 1 |
| adductor longus | 7.51 ± 29.04 | -7.51 ± 23.27 | 36.2 | -2.41 | 0.678 |
| adductor magnus | 13.06 ± 153.55 | -13.06 ± 97.56 | 36.1 | -0.81 | 1 |
| anconeus | 0.69 ± 2.1 | -0.73 ± 1.67 | 36.8 | -3.26 | 0.111 |
| biceps brachii | 14.75 ± 40.13 | -16.08 ± 24.11 | 31 | -3.66 | 0.053 |
| biceps femoris: long head | 9.19 ± 36.95 | -9.19 ± 32 | 38.5 | -2.25 | 0.89 |
| biceps femoris: short head | 5.88 ± 16.76 | -5.88 ± 14.75 | 38.5 | -2.89 | 0.259 |
| brachialis | 7.44 ± 25.16 | -8.14 ± 18.94 | 34.8 | -2.85 | 0.28 |
| brachioradialis | 7.07 ± 16.23 | -7.69 ± 10.73 | 35.6 | -4.12 | 0.013 |
| coracobrachialis | 1.89 ± 6.21 | -2.06 ± 5.88 | 38.5 | -3.21 | 0.122 |
| deep forearm extensors | 2.08 ± 5.62 | -2.19 ± 4.75 | 38.8 | -3.57 | 0.055 |
| deltoid | 27.94 ± 79.29 | -29.86 ± 53.06 | 30.4 | -3.55 | 0.067 |
| digit extensors | 1.98 ± 8.41 | -2.02 ± 4.74 | 32.1 | -1.93 | 1 |
| erector spinae ub | 23.94 ± 85.3 | -25.09 ± 63.83 | 38.5 | -1.7 | 1 |
| extensor carpi radialis | 4.52 ± 11.08 | -4.91 ± 8.01 | 36.9 | -4.16 | 0.012 |
| extensor carpi ulnaris | 0.85 ± 5 | -0.73 ± 2.93 | 33.6 | -1.39 | 1 |
| external oblique | 4.55 ± 27.6 | -4.45 ± 40.11 | 27.1 | -0.61 | 1 |
| fibulari | 0.18 ± 19.42 | -0.18 ± 19.47 | 33.9 | -0.24 | 1 |
| flexor carpi radialis and palmaris longus | 3.09 ± 12.03 | -3.23 ± 7.21 | 35.8 | -2.46 | 0.62 |
| flexor carpi ulnaris | 2.15 ± 8.12 | -2.46 ± 5.72 | 35.5 | -3.33 | 0.1 |
| flexor digitorum longus | 0.99 ± 4.77 | -0.99 ± 3.34 | 39 | -1.43 | 1 |
| flexor digitorum profundus | 5.48 ± 15.07 | -6.13 ± 11.76 | 38.3 | -3.54 | 0.059 |
| flexor digitorum superficialis | 5.67 ± 12.83 | -6.07 ± 8.2 | 35 | -4.27 | 0.009 |
| flexor hallucis longus | 3.09 ± 11.42 | -3.09 ± 9.22 | 39 | -1.82 | 1 |
| flexor pollicis longus | 1.09 ± 3.82 | -1.15 ± 2.75 | 32.7 | -2.71 | 0.368 |
| gastrocnemius: lateral head | -1.42 ± 24.57 | 1.42 ± 28.74 | 35.4 | 0.49 | 1 |
| gastrocnemius: medial head | -2.58 ± 52.36 | 2.58 ± 38.16 | 37.6 | 0.57 | 1 |
| gemelli | 0.44 ± 3.97 | -0.44 ± 3.49 | 39 | -1.34 | 1 |
| gluteus maximus | 15.55 ± 159.1 | -15.55 ± 100.25 | 36.7 | -0.8 | 1 |
| gluteus medius | 12.95 ± 47.47 | -12.95 ± 34.11 | 36 | -2.78 | 0.319 |
| gluteus minimus | 3.55 ± 15.88 | -3.55 ± 10.21 | 32.5 | -1.9 | 1 |
| gracilis | 3.06 ± 21.73 | -2.93 ± 13.94 | 35.6 | -1.62 | 1 |
| iliacus | 12.75 ± 21.34 | -12.75 ± 21.52 | 38.5 | -4.54 | 0.004 |
| infraspinatus and teres minor | 12.62 ± 26.68 | -13.56 ± 24.41 | 37 | -4.21 | 0.01 |
| latissimus dorsi | 26.14 ± 89.7 | -31.92 ± 65.83 | 37.7 | -2.26 | 0.89 |
| levator scapulae | 4.19 ± 9.11 | -4.66 ± 8.28 | 34.3 | -4.33 | 0.008 |
| multifidus ub | 6.02 ± 27.08 | -7.44 ± 18.86 | 36 | -1.96 | 1 |
| obturator externus | 1.52 ± 10.77 | -1.52 ± 6.44 | 32.8 | -1.56 | 1 |
| obturator internus | 1.49 ± 5.27 | -1.49 ± 3.87 | 37.1 | -2.85 | 0.28 |
| pectineus | 3.5 ± 10.74 | -3.5 ± 7.37 | 30.9 | -3.51 | 0.071 |
| pectoralis major | 37.03 ± 85.32 | -39.4 ± 60.13 | 34.1 | -3.86 | 0.029 |
| pectoralis minor | 3.72 ± 8.16 | -3.84 ± 8.11 | 38.3 | -3.64 | 0.046 |
| phalangeal extensors | 0.62 ± 9.96 | -0.62 ± 13.83 | 29.7 | -0.03 | 1 |
| piriformis | 1.05 ± 8.33 | -1.05 ± 8.96 | 36.8 | -0.83 | 1 |
| popliteus | 1.19 ± 3.65 | -1.33 ± 2.31 | 33.5 | -2.74 | 0.355 |
| pronator quadratus | 0.65 ± 2.21 | -0.77 ± 1.65 | 38.6 | -2.99 | 0.21 |
| pronator teres | 1.76 ± 7.18 | -2.03 ± 5.97 | 38.8 | -2.54 | 0.521 |
| psoas major | 24.2 ± 46.14 | -25.25 ± 32.96 | 35.9 | -4.57 | 0.004 |
| quadratus femoris | 2.32 ± 7.35 | -2.43 ± 6.3 | 38.5 | -2.87 | 0.266 |
| quadratus lumborum | 4.35 ± 11.35 | -4.54 ± 8.21 | 38.9 | -3.01 | 0.203 |
| rectus abdominis | -3.4 ± 45.58 | 3.55 ± 48.87 | 36.5 | 0.72 | 1 |
| rectus femoris | 11.36 ± 46.92 | -11.36 ± 37.85 | 38.1 | -2.31 | 0.82 |
| rhomboid | 6.64 ± 12.65 | -7.36 ± 10.01 | 36 | -5.23 | 0.001 |
| sartorius | 6.38 ± 24.32 | -6.38 ± 23.06 | 38.3 | -1.62 | 1 |
| semimembranosus | 6.32 ± 43.18 | -6.32 ± 35.07 | 38.9 | -0.87 | 1 |
| semitendinosus | 5.26 ± 37.47 | -5.04 ± 25.28 | 37.3 | -1.28 | 1 |
| serratus anterior | 12.23 ± 26.98 | -13.41 ± 21.19 | 37.6 | -4.37 | 0.006 |
| soleus | 1.09 ± 57.14 | -1.09 ± 48.73 | 39 | 0.04 | 1 |
| subclavius | 0.72 ± 2.5 | -0.78 ± 1.23 | 27.5 | -3.57 | 0.07 |

| | | | | | |
|---|---|---|---|---|---|
| **subscapularis** | 10 ± 21.57 | -10.64 ± 20.07 | 36.6 | -5.47 | < 0.001 |
| **supinator** | 1.09 ± 4.13 | -1.3 ± 2.92 | 30.9 | -3.96 | 0.025 |
| **supraspinatus** | 5.01 ± 11.85 | -5.46 ± 10.49 | 38.9 | -3.53 | 0.059 |
| **tensor fasciae latae** | 4.51 ± 19.6 | -4.51 ± 11.74 | 37.8 | -1.55 | 1 |
| **teres major** | 2.46 ± 7.14 | -2.56 ± 4.96 | 35.5 | -3.37 | 0.092 |
| **tibialis anterior** | 1.95 ± 12.21 | -1.95 ± 12.42 | 37.1 | -1.14 | 1 |
| **tibialis posterior** | 3.99 ± 15.57 | -3.99 ± 14.82 | 36 | -1.13 | 1 |
| **trapezius** | 15.56 ± 40.18 | -18.11 ± 28.89 | 37.2 | -3.11 | 0.162 |
| **triceps brachii** | 28.12 ± 96.25 | -30.57 ± 64.99 | 32 | -3.02 | 0.21 |
| **vastus intermedius** | 12.03 ± 59.74 | -12.03 ± 30.99 | 31.2 | -1.56 | 1 |
| **vastus lateralis** | 66.17 ± 162.84 | -66.17 ± 107.48 | 32.4 | -3.71 | 0.046 |
| **vastus medialis** | 31.53 ± 84.61 | -31.53 ± 56.29 | 34.7 | -3.29 | 0.11 |

**Table S8:** Post-hoc analysis results of the interaction effect for the dependent variable of actual versus expected volume difference using the associated bone volume linear regression model reported for individual muscles. The contrast was Biologic Sex*Muscle.

| Muscle/Group Names | Mean ± STDEV (mL) | | DoF | t-Statistic | p-Value |
|---|---|---|---|---|---|
| | Male | Female | | | |
| adductor brevis | -0.35 ± 16.32 | 0.35 ± 14.91 | 38.6 | 0.5 | 1 |
| adductor longus | 5.8 ± 29.3 | -5.8 ± 28.2 | 38.8 | -1.11 | 1 |
| adductor magnus | 1.88 ± 149.25 | -1.88 ± 110.09 | 38.4 | 0.23 | 1 |
| anconeus | 0.22 ± 1.92 | -0.15 ± 1.55 | 38.1 | -0.78 | 1 |
| biceps brachii | 5.79 ± 36.28 | -5.36 ± 25.08 | 37.4 | -1.08 | 1 |
| biceps femoris: long head | 6.4 ± 34.22 | -6.4 ± 36.5 | 36.8 | -1.13 | 1 |
| biceps femoris: short head | 5.03 ± 18.73 | -5.03 ± 16.72 | 38.3 | -1.33 | 1 |
| brachialis | 2.03 ± 23.51 | -1.62 ± 18.11 | 38.1 | -0.5 | 1 |
| brachioradialis | 3.71 ± 15.48 | -3.67 ± 11.56 | 38.6 | -1.69 | 1 |
| coracobrachialis | 0.74 ± 5.54 | -0.68 ± 6.3 | 33.3 | -1.11 | 1 |
| deep forearm extensors | 0.73 ± 4.98 | -0.51 ± 4.81 | 38.7 | -1.25 | 1 |
| deltoid | 9.25 ± 63.03 | -7.34 ± 61.89 | 38.4 | -0.77 | 1 |
| digit extensors | -0.07 ± 7.34 | 0.33 ± 4.98 | 38.2 | 0.81 | 1 |
| erector spinae ub | 32.27 ± 69.46 | -31.6 ± 90.25 | 31.6 | -1.52 | 1 |
| extensor carpi radialis | 2.01 ± 11.09 | -1.66 ± 7.28 | 37.2 | -1.4 | 1 |
| extensor carpi ulnaris | 0.01 ± 5.01 | 0.27 ± 3.02 | 36.1 | 0.82 | 1 |
| external oblique | 10.76 ± 24.72 | -10.54 ± 50.18 | 22.3 | -1.08 | 1 |
| fibulari | 0.79 ± 19.23 | -0.79 ± 21.8 | 31.1 | -0.15 | 1 |
| flexor carpi radialis and palmaris longus | 1.05 ± 11.15 | -0.63 ± 8.01 | 38.3 | -0.45 | 1 |
| flexor carpi ulnaris | 0.57 ± 7.68 | -0.43 ± 5.69 | 37.9 | -0.88 | 1 |
| flexor digitorum longus | 1.32 ± 5.46 | -1.32 ± 3.86 | 38.9 | -1.3 | 1 |
| flexor digitorum profundus | 2.12 ± 14.36 | -1.81 ± 12.13 | 38.9 | -1.06 | 1 |
| flexor digitorum superficialis | 2.27 ± 10.83 | -1.88 ± 8.28 | 38.4 | -1.62 | 1 |
| flexor hallucis longus | 3.33 ± 12.37 | -3.33 ± 9.91 | 39 | -1.44 | 1 |
| flexor pollicis longus | 0.3 ± 3.28 | -0.15 ± 2.72 | 38.5 | -0.44 | 1 |
| gastrocnemius: lateral head | 0.8 ± 24.18 | -0.8 ± 34.49 | 30.7 | 0.23 | 1 |
| gastrocnemius: medial head | -2.66 ± 49.81 | 2.66 ± 42.93 | 39 | 0.77 | 1 |
| gemelli | 0.3 ± 3.69 | -0.3 ± 3.03 | 38.9 | -0.75 | 1 |
| gluteus maximus | 25.01 ± 130.54 | -25.01 ± 152.27 | 36.4 | -0.61 | 1 |
| gluteus medius | 14.69 ± 38.85 | -14.69 ± 43.91 | 34.9 | -2.27 | 1 |
| gluteus minimus | 3.14 ± 11.44 | -3.14 ± 11.68 | 37.1 | -1.26 | 1 |
| gracilis | 2.03 ± 21.93 | -1.95 ± 18.81 | 38.8 | -0.46 | 1 |
| iliacus | 14.93 ± 23.24 | -14.93 ± 23.27 | 37.9 | -3.91 | 0.026 |
| infraspinatus and teres minor | 5.55 ± 24.92 | -5.1 ± 23.13 | 39 | -1.43 | 1 |
| latissimus dorsi | 9.55 ± 74.92 | -9.1 ± 83.59 | 36.4 | -0.32 | 1 |
| levator scapulae | 2.14 ± 8.61 | -2.05 ± 7.78 | 36 | -2.04 | 1 |
| multifidus ub | 8.93 ± 24.73 | -8.54 ± 28.3 | 33.1 | -1.61 | 1 |
| obturator externus | 1.14 ± 9.51 | -1.14 ± 5.73 | 34.3 | -1 | 1 |
| obturator internus | 1 ± 4.25 | -1 ± 3.64 | 39 | -2.21 | 1 |
| pectineus | 4.51 ± 10.88 | -4.51 ± 9.74 | 39 | -2.83 | 0.506 |
| pectoralis major | 15.98 ± 70.26 | -14.75 ± 62.2 | 38.9 | -1.37 | 1 |
| pectoralis minor | 2.53 ± 9.35 | -2.25 ± 8.53 | 38.2 | -1.65 | 1 |
| phalangeal extensors | 0.65 ± 9.81 | -0.65 ± 14.74 | 29.2 | 0.26 | 1 |
| piriformis | 0.77 ± 7.69 | -0.77 ± 9.16 | 34.2 | -0.48 | 1 |
| popliteus | 0.83 ± 3.54 | -0.89 ± 2.63 | 37.5 | -1.48 | 1 |
| pronator quadratus | 0.32 ± 2.2 | -0.36 ± 1.6 | 37.3 | -1.51 | 1 |
| pronator teres | 0.72 ± 7.3 | -0.57 ± 6.52 | 38.6 | -0.48 | 1 |
| psoas major | 26.3 ± 41.2 | -27.44 ± 41.32 | 38.2 | -4.1 | 0.015 |
| quadratus femoris | 1.7 ± 6.47 | -1.78 ± 5.78 | 38.9 | -2.08 | 1 |
| quadratus lumborum | 4.79 ± 10.16 | -5 ± 10.14 | 35.9 | -2.8 | 0.558 |
| rectus abdominis | 2.12 ± 43.1 | -2.21 ± 63.18 | 30.7 | 0.24 | 1 |
| rectus femoris | 3.29 ± 39.84 | -3.29 ± 36.62 | 38.8 | -0.51 | 1 |
| rhomboid | 3.58 ± 13.22 | -3.62 ± 10.57 | 38.6 | -1.97 | 1 |
| sartorius | 7.41 ± 28.78 | -7.41 ± 31.05 | 36.5 | -0.92 | 1 |
| semimembranosus | 2.59 ± 40.37 | -2.59 ± 39.91 | 37.6 | 0.27 | 1 |
| semitendinosus | 2.81 ± 33.76 | -2.7 ± 37.01 | 36.4 | -0.09 | 1 |
| serratus anterior | 5.57 ± 22.36 | -5.2 ± 24.24 | 34.1 | -1.61 | 1 |
| soleus | 4.27 ± 58.12 | -4.27 ± 59.64 | 38.3 | -0.01 | 1 |
| subclavius | 0.51 ± 2.6 | -0.52 ± 1.34 | 30.9 | -2.12 | 1 |

| | | | | | |
|---|---|---|---|---|---|
| **subscapularis** | 4.01 ± 19.39 | -3.59 ± 17.64 | 38.9 | -2.03 | 1 |
| **supinator** | 0.25 ± 3.74 | -0.26 ± 2.86 | 37.3 | -1.09 | 1 |
| **supraspinatus** | 1.63 ± 10.13 | -1.56 ± 8.73 | 39 | -1.05 | 1 |
| **tensor fasciae latae** | 5.48 ± 17.25 | -5.48 ± 16.2 | 34.9 | -1.41 | 1 |
| **teres major** | 1.23 ± 6.86 | -1.12 ± 4.69 | 36.6 | -1.58 | 1 |
| **tibialis anterior** | 1.95 ± 12.9 | -1.95 ± 13.15 | 36.4 | -0.65 | 1 |
| **tibialis posterior** | 4.44 ± 16.86 | -4.44 ± 14.5 | 38.5 | -1.07 | 1 |
| **trapezius** | 7.11 ± 29.7 | -6.32 ± 36.31 | 32.7 | -0.61 | 1 |
| **triceps brachii** | 7.35 ± 82.56 | -5.86 ± 66.64 | 38.5 | -0.64 | 1 |
| **vastus intermedius** | 8.63 ± 59.19 | -8.63 ± 42.26 | 36.7 | -0.44 | 1 |
| **vastus lateralis** | 36.71 ± 143.6 | -36.71 ± 101.76 | 36.5 | -1.7 | 1 |
| **vastus medialis** | 16.27 ± 75.86 | -16.27 ± 51.42 | 36 | -1.3 | 1 |

**Table S9:** Linear Regression Model Elements of the Associated Bone Volume Scaling Method for Male and Female Cohorts

| Muscle/Group Name | Combined | | Male | | Female | |
|---|---|---|---|---|---|---|
| | Slope | Intercept | Slope | Intercept | Slope | Intercept |
| Levator Scapulae | 0.272 | 4.06 | 0.162 | 26.048 | 0.168 | 14.698 |
| Supraspinatus | 0.431 | -0.518 | 0.32 | 21.115 | 0.444 | 3.892 |
| Trapezius | 1.482 | -0.074 | 1.064 | 82.772 | 1.329 | 11.381 |
| Rhomboid | 0.534 | -9.594 | 0.354 | 26.629 | 0.351 | 9.487 |
| Serratus Anterior | 1.096 | 5.032 | 0.937 | 39.534 | 0.398 | 86.192 |
| Latissimus Dorsi | 2.959 | -34.093 | 2.571 | 45.739 | 2.127 | 60.006 |
| Subscapularis | 0.857 | 3.321 | 0.701 | 35.502 | 0.492 | 44.62 |
| Teres Major | 0.173 | 0.917 | 0.104 | 14.551 | 0.131 | 4.831 |
| Infraspinatus & Teres Minor | 1.101 | -0.067 | 0.87 | 47.295 | 0.651 | 50.281 |
| Subclavius | 0.044 | 0.051 | 0.02 | 5.063 | 0.014 | 3.272 |
| Pectoralis Minor | 0.29 | 2.072 | 0.157 | 28.775 | 0.182 | 12.981 |
| Pectoralis Major | 2.932 | -109.18 | 2.197 | 40.027 | 1.877 | 5.78 |
| Deltoid | 3.027 | -46.08 | 2.832 | 1.458 | 1.629 | 118.602 |
| Triceps Brachii | 3.156 | -72.743 | 3.035 | 43.474 | 1.928 | 72.672 |
| Coracobrachialis | 0.187 | 1.263 | 0.165 | 6.071 | 0.099 | 11.477 |
| Biceps Brachii | 1.404 | -51.401 | 1.253 | 18.158 | 0.629 | 39.271 |
| Brachialis | 0.879 | -1.846 | 0.884 | 0.786 | 0.407 | 54.809 |
| Brachioradialis | 0.533 | -15.974 | 0.396 | 12.469 | 0.17 | 25.4 |
| Extensor Carpi Radialis | 1.267 | 5.403 | 0.928 | 24.434 | 0.86 | 16.663 |
| Anconeus | 0.22 | 0.414 | 0.196 | 1.843 | 0.137 | 2.909 |
| Digit Extensors | 0.876 | 3.578 | 0.931 | 0.817 | 0.702 | 9.288 |
| Extensor Carpi Ulnaris | 0.436 | 1.191 | 0.451 | 0.381 | 0.396 | 2.512 |
| Deep Forearm Extensors | 0.63 | 5.734 | 0.549 | 10.499 | 0.336 | 14.645 |
| Supinator | 0.352 | 1.599 | 0.334 | 2.799 | 0.222 | 5.626 |
| Pronator Teres | 0.703 | 6.786 | 0.585 | 13.453 | 0.546 | 11.206 |
| Flexor Carpi Radialis & Palmaris Longus | 1.06 | 8.463 | 0.867 | 19.086 | 0.867 | 13.711 |
| Flexor Digitorum Superficialis | 1.481 | 2.909 | 1.06 | 26.174 | 1.094 | 13.325 |
| Flexor Carpi Ulnaris | 0.837 | 5.01 | 0.853 | 4.761 | 0.342 | 20.771 |
| Flexor Digitorum Profundus | 1.794 | 10.977 | 1.445 | 30.5 | 1.277 | 25.82 |
| Flexor Pollicis Longus | 0.349 | 2.009 | 0.308 | 4.388 | 0.261 | 4.591 |
| Pronator Quadratus | 0.147 | 1.877 | 0.084 | 5.307 | 0.103 | 2.975 |
| Rectus Abdominis | 0.378 | 55.379 | 0.38 | 56.316 | 0.303 | 88.051 |
| External Oblique | 0.244 | 55.21 | 0.193 | 95.435 | 0.012 | 153.204 |
| Erector Spinae | 1.053 | -72.71 | 0.789 | 112.106 | 0.554 | 127.826 |
| Multifidus | 0.377 | 4.311 | 0.294 | 61.531 | 0.258 | 50.681 |
| Quadratus Lumborum | 0.107 | 2.887 | 0.043 | 44.362 | 0.079 | 10.83 |
| Psoas Major | 0.589 | -50.289 | 0.35 | 114.119 | 0.211 | 98.23 |
| Iliacus | 0.347 | -9.199 | 0.226 | 75.488 | 0.111 | 85.925 |
| Gluteus Medius | 0.59 | 38.065 | 0.526 | 89.915 | 0.248 | 183.114 |
| Gluteus Maximus | 1.972 | -77.639 | 1.936 | 31.668 | 1.242 | 238.577 |
| Gluteus Minimus | 0.193 | -1.642 | 0.18 | 9.152 | 0.119 | 29.818 |
| Piriformis | 0.038 | 22.82 | 0.031 | 27.827 | 0.028 | 26.727 |
| Gemelli | 0.041 | -3.388 | 0.043 | 4.058 | 0.028 | 2.429 |
| Quadratus Femoris | 0.061 | -4.872 | 0.054 | 0.934 | 0.011 | 16.823 |
| Obturator Internus | 0.045 | -2.724 | 0.038 | 2.128 | 0.026 | 4.916 |
| Obturator Externus | 0.104 | -7.987 | 0.097 | 3.055 | 0.081 | 1.779 |
| Pectineus | 0.156 | -21.635 | 0.135 | 4.901 | 0.054 | 21.5 |
| Tensor Fasciae Latae | 0.171 | -16.445 | 0.148 | 2.057 | 0.04 | 38.984 |
| Rectus Femoris | 0.606 | -24.975 | 0.595 | 15.81 | 0.416 | 46.213 |
| Vastus Lateralis | 2.081 | -41.916 | 1.58 | 262.332 | 1.207 | 265.001 |
| Vastus Intermedius | 0.566 | -3.005 | 0.428 | 79.685 | 0.429 | 42.281 |
| Vastus Medialis | 1.137 | -53.219 | 0.913 | 82.55 | 0.755 | 80.628 |
| Sartorius | 0.284 | 20.977 | 0.153 | 98.358 | 0.205 | 44.445 |
| Adductor Brevis | 0.183 | 14.026 | 0.209 | 0.086 | 0.123 | 37.968 |
| Adductor Magnus | 1.365 | -22.771 | 1.412 | 45.598 | 1.086 | 85.306 |
| Adductor Longus | 0.313 | 23.756 | 0.245 | 65.636 | 0.136 | 87.218 |
| Gracilis | 0.229 | -4.935 | 0.214 | 4.936 | 0.144 | 26.305 |
| Semitendinosus | 0.437 | -9.723 | 0.418 | 3.271 | 0.316 | 35.137 |
| Semimembranosus | 0.438 | 38.514 | 0.413 | 54.6 | 0.344 | 73.084 |
| Biceps Femoris: Long Head | 0.324 | 54.983 | 0.209 | 122.97 | 0.264 | 72.439 |
| Biceps Femoris: Short Head | 0.194 | 3.739 | 0.131 | 42.396 | 0.056 | 53.143 |
| Popliteus | 0.032 | 3.559 | 0.015 | 13.519 | 0.031 | 3.06 |
| Gastrocnemius: Medial Head | 0.518 | 107.753 | 0.533 | 100.289 | 0.663 | 75.018 |
| Gastrocnemius: Lateral Head | 0.303 | 60.211 | 0.284 | 67.075 | 0.296 | 61.147 |
| Soleus | 0.765 | 218.634 | 0.521 | 301.269 | 1.094 | 134.306 |
| Tibialis Anterior | 0.261 | 38.586 | 0.233 | 49.696 | 0.2 | 51.646 |
| Phalangeal Extensors | 0.198 | 30.713 | 0.154 | 45.668 | 0.267 | 13.344 |
| Fibulari | 0.239 | 53.855 | 0.172 | 76.343 | 0.358 | 24.251 |

| | | | | | | |
|---|---|---|---|---|---|---|
| Tibialis Posterior | 0.15 | 47.111 | 0.013 | 95.474 | 0.192 | 32.316 |
| Flexor Digitorum Longus | 0.071 | 2.457 | 0.045 | 12.267 | 0.048 | 6.805 |
| Flexor Hallucis Longus | 0.193 | 13.322 | 0.109 | 43.711 | 0.178 | 13.59 |

**Atlas:**

**Figure S1A:** Upper Body Atlas. The following muscles are outlined in the axial image slices and labeled accordingly: Multifidus (MF), Erector Spinae (ES), Rhomboid (RH), Trapezius (TR), Levator Scapulae (LS), Serratus Anterior (SA), Pectoralis Major (PMA), Pectoralis Minor (PMI), Deltoid (DE), Coracobrachialis (CB), Biceps Brachii (BB), Triceps Brachii (TB), Latissimus Dorsi (LD), Subscapularis (SC), Teres Major (TM), Infraspinatus/Teres Minor (ITM), Brachialis (BC), Brachioradialis (BR), Extensor Carpi Radialis (ECR), Flexor Carpi Radialis & Palmaris Longus (FCR/PL), Flexor Digitorum Superficialis (FDS), Flexor Carpi Ulnaris (FCU), Flexor Digitorum Profundus (FDP), Flexor Pollicis Longus (FPL), Deep Forearm Extensors (DFE), Digit Extensors (DE), Extensor Carpi Ulnaris (ECU).

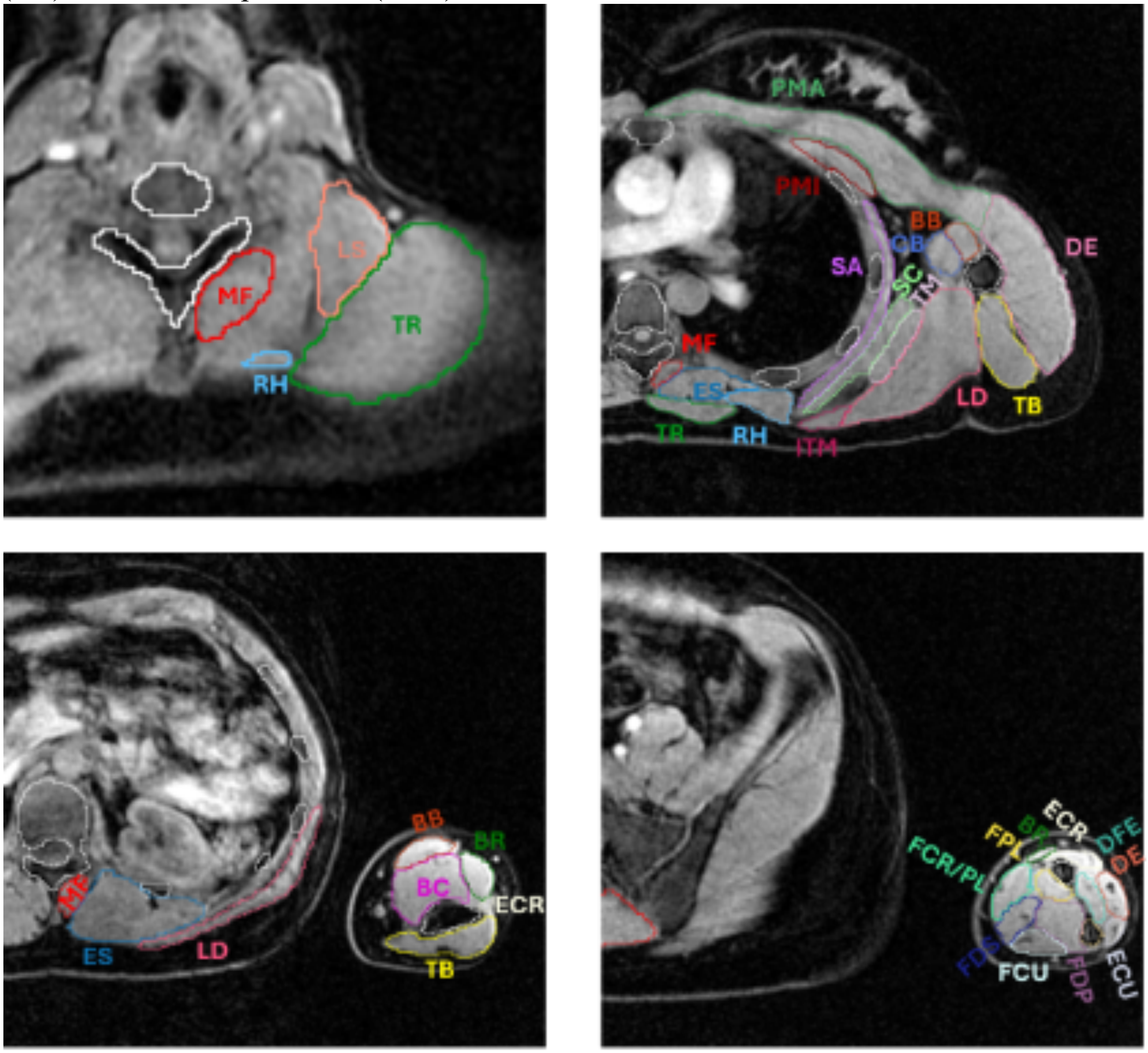


**Figure S1B:** Lower Extremity Atlas. The following muscles are outlined in the axial image slices and labeled accordingly: Rectus Abdominis (RA), Extenal Oblique (EO), Multifidus (MF), Erector Spinae (ES), Psoas Major (PM), Quadratus Lumborum (QL), Gluteus Maximus (GMAX), Quadratus Femoris (QF), Adductor Magnus (AM), Adductor Brevis (AB), Adductor Longus (AL), Pectineus (PE), Iliacus (IL), Vastus Intermedius (VI), Vastus Lateralis (VL), Tensor Fasciae Latae

(TFL), Rectus Femoris (RF), Sartorius (SA), Vastus Medialis (VM), Semimembranosus (SM), Semitendinosus (ST), Biceps Femoris Long Head (BFLH), Biceps Femoris Short Head (BFSH), Gracilis (GR), Tibialis Anterior (TA), Fibulari (FI), Phalangeal Extensors (PE), Tibialis Posterior (TP), Soleus (SO), Gastrocnemius Lateral Head (GLH), Gastrocnemius Medial Head (GMH).

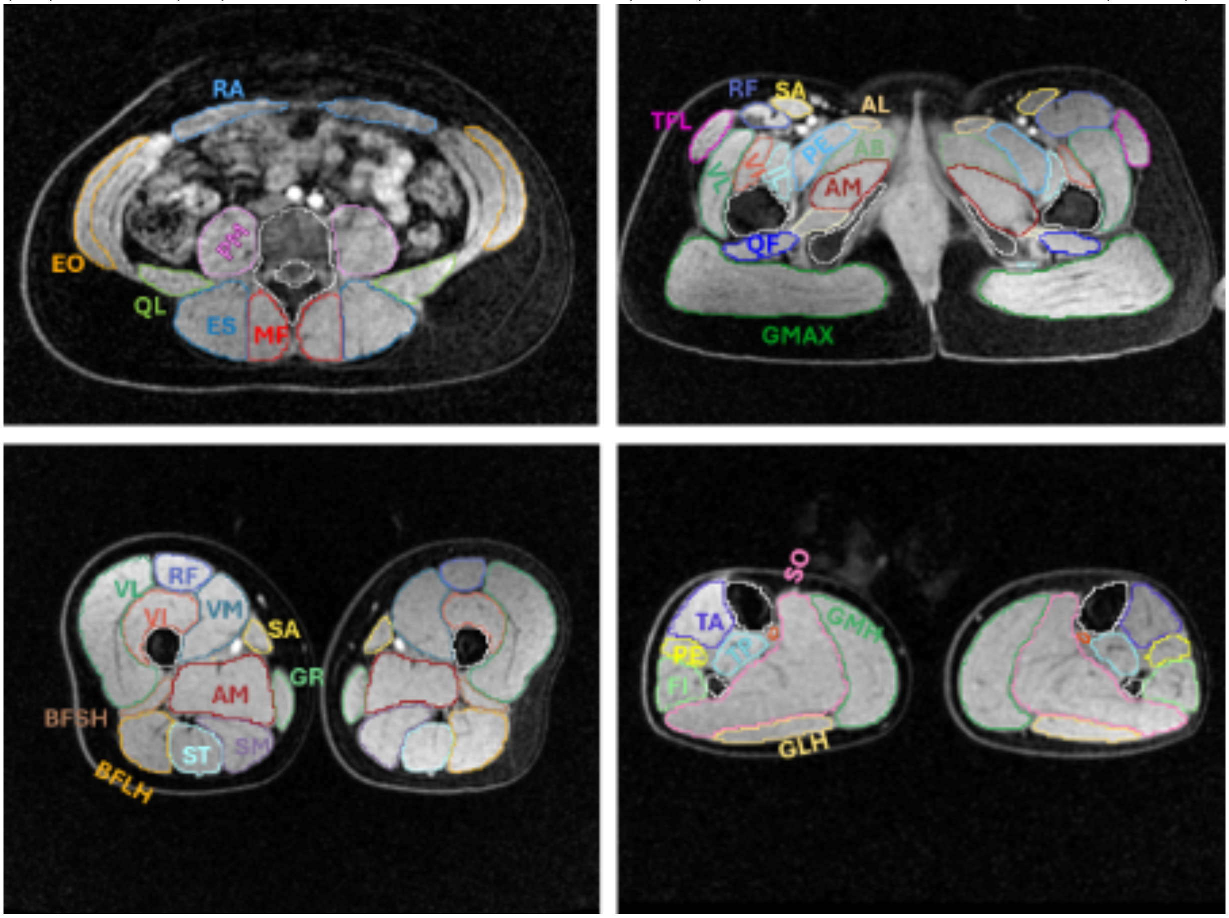

### Muscle Volume Comparisons to Literature

Average volume fractions were calculated as a percentage of the total left/right muscle region volume. Individual left and right muscles were calculated separately, resulting in both a left and right volume fraction for each muscle. Total left and right muscle region volumes were also calculated separately by summing the volumes of individual muscles from each side. To directly compare the upper body (UB) muscle volume fractions to those reported in literature, a subset of the UB muscles reported in this study were used to calculate the total left and right upper body muscle region volume. The same method was used to find the average volume fractions for the Lower Extremity (LE) muscles.

**Table S10:** Average muscle volume fractions found in this study compared to muscle volume fractions found in literature [1,2].

| Muscle | Muscle Region | Average Volume Fraction (%) ± SD | Literature Average Volume Fraction (%) ± SD |
|---|---|---|---|
| latissimus dorsi | UB | 14.68 ± 1.16 | 9.8 ± 1.52 |
| pectoralis major | UB | 11.69 ± 1.22 | 10.72 ± 1.97 |
| deltoid | UB | 14.65 ± 0.75 | 15.2 ± 1.05 |
| supraspinatus | UB | 2.33 ± 0.34 | 2.02 ± 0.32 |
| infraspinatus and teres minor | UB | 5.94 ± 0.52 | 5.9 ± 0.75 |
| subscapularis | UB | 4.77 ± 0.61 | 6.64 ± 0.78 |
| teres major | UB | 0.96 ± 0.17 | 1.28 ± 0.25 |
| coracobrachialis | UB | 1.06 ± 0.18 | 0.93 ± 0.28 |
| triceps | UB | 14.28 ± 0.97 | 14.53 ± 0.69 |
| biceps | UB | 5.66 ± 0.5 | 5.62 ± 0.46 |
| brachialis | UB | 4.7 ± 0.5 | 5.69 ± 0.67 |
| brachioradialis | UB | 2.28 ± 0.31 | 2.49 ± 0.47 |
| anconeus | UB | 0.34 ± 0.06 | 0.43 ± 0.08 |
| supinator | UB | 0.58 ± 0.11 | 0.83 ± 0.23 |
| pronator teres | UB | 1.28 ± 0.2 | 1.52 ± 0.25 |
| pronator quadratus | UB | 0.29 ± 0.07 | 0.44 ± 0.11 |
| extensor carpi radialis | UB | 2.05 ± 0.19 | 2.33 ± 0.22 |
| extensor carpi ulnaris | UB | 0.69 ± 0.13 | 0.67 ± 0.09 |
| flexor carpi radialis and palmaris longus | UB | 1.87 ± 0.27 | 1.52 ± 0.36 |

| | | | |
|---|---|---|---|
| flexor carpi ulnaris | UB | 1.42 ± 0.19 | 1.54 ± 0.33 |
| digit extensors | UB | 1.43 ± 0.22 | 1.42 ± 0.16 |
| deep forearm extensors | UB | 1.15 ± 0.19 | 1.08 ± 0.11 |
| flexor digitorum superficialis | UB | 2.28 ± 0.24 | 3.05 ± 0.45 |
| flexor digitorum profundus | UB | 3.04 ± 0.34 | 3.65 ± 0.45 |
| flexor pollicis longus | UB | 0.59 ± 0.09 | 0.71 ± 0.16 |
| gluteus maximus | LE | 13.17 ± 1.23 | 11.93 ± 1.02 |
| adductor magnus | LE | 8.37 ± 1.08 | 7.86 ± 0.86 |
| gluteus medius | LE | 4.83 ± 0.46 | 4.54 ± 0.62 |
| psoas major | LE | 3.51 ± 0.57 | 3.8 ± 0.61 |
| iliacus | LE | 2.36 ± 0.29 | 2.48 ± 0.31 |
| sartorius | LE | 2.11 ± 0.33 | 2.29 ± 0.29 |
| adductor longus | LE | 2.34 ± 0.28 | 2.26 ± 0.34 |
| gluteus minimus | LE | 1.39 ± 0.17 | 1.47 ± 0.19 |
| adductor brevis | LE | 1.37 ± 0.17 | 1.47 ± 0.23 |
| gracilis | LE | 1.38 ± 0.2 | 1.46 ± 0.23 |
| pectineus | LE | 0.82 ± 0.12 | 0.92 ± 0.22 |
| tensor fasciae latae | LE | 0.99 ± 0.2 | 0.89 ± 0.27 |
| obturator externus | LE | 0.64 ± 0.1 | 0.76 ± 0.16 |
| piriformis | LE | 0.61 ± 0.15 | 0.61 ± 0.17 |
| quadratus femoris | LE | 0.37 ± 0.08 | 0.45 ± 0.11 |
| obturator internus | LE | 0.29 ± 0.06 | 0.38 ± 0.09 |
| gemelli | LE | 0.25 ± 0.05 | 0.23 ± 0.07 |
| vastus lateralis | LE | 12.6 ± 1.31 | 11.66 ± 1.06 |
| vastus medialis | LE | 6.49 ± 0.6 | 6.06 ± 0.56 |
| vastus intermedius | LE | 3.57 ± 0.42 | 3.84 ± 0.6 |
| rectus femoris | LE | 3.51 ± 0.45 | 3.79 ± 0.51 |
| semimembranosus | LE | 3.35 ± 0.4 | 3.46 ± 0.43 |
| biceps femoris: long head | LE | 2.83 ± 0.36 | 2.92 ± 0.43 |
| semitendinosus | LE | 2.64 ± 0.32 | 2.6 ± 0.34 |
| biceps femoris: short head | LE | 1.28 ± 0.19 | 1.4 ± 0.3 |
| popliteus | LE | 0.26 ± 0.04 | 0.33 ± 0.04 |
| soleus | LE | 6.1 ± 0.82 | 6.21 ± 0.8 |
| gastrocnemius: medial head | LE | 3.56 ± 0.61 | 3.62 ± 0.48 |
| gastrocnemius: lateral head | LE | 2.05 ± 0.36 | 2.11 ± 0.42 |
| tibialis anterior | LE | 1.56 ± 0.18 | 1.91 ± 0.23 |
| fibulari | LE | 1.69 ± 0.27 | 1.83 ± 0.28 |
| tibialis posterior | LE | 1.26 ± 0.25 | 1.49 ± 0.24 |

| | | | |
|---|---|---|---|
| phalangeal extensors | LE | 1.21 ± 0.16 | 1.44 ± 0.14 |
| flexor hallucis longus | LE | 0.94 ± 0.13 | 1.09 ± 0.19 |
| flexor digitorum longus | LE | 0.31 ± 0.06 | 0.43 ± 0.11 |


1 Handsfield, G. G., Meyer, C. H., Hart, J. M., Abel, M. F. & Blemker, S. S. Relationships of 35 lower limb muscles to height and body mass quantified using MRI. *J Biomech* **47**, 631-638 (2014). https://doi.org:10.1016/j.jbiomech.2013.12.002

2 Holzbaur, K. R., Murray, W. M., Gold, G. E. & Delp, S. L. Upper limb muscle volumes in adult subjects. *J Biomech* **40**, 742-749 (2007). https://doi.org:10.1016/j.jbiomech.2006.11.011
